# Striatal endocannabinoid-long-term potentiation mediates one-shot learning

**DOI:** 10.1101/2024.07.10.602953

**Authors:** Charlotte Piette, Arnaud Hubert, Sylvie Perez, Hugues Berry, Jonathan Touboul, Laurent Venance

## Abstract

One-shot learning, the ability to form memories after a single, brief salient event, is essential to adapt one’s behaviour in a dynamic world. However, how one-shot learning unfolds in the brain remains unknown. Challenges to elucidate its neural underpinnings stem experimentally from the scarcity of behavioral assays recapitulating one-shot learning in the laboratory, and conceptually from the limited number of neuronal plasticity mechanisms that could support learning after a small number of action potentials, as is common during a one-shot experience. Here, we overcome these challenges and identify a new mechanism for one-shot learning in dorsal striatum that relies on a non-classical form of plasticity, the endocannabinoid-mediated long-term potentiation (eCB-LTP). To do so, we develop a novel one-shot behavioral test, in which mice learn to avoid a sticky tape after a single, spontaneous and brief contact with the uncomfortable substrate, and maintain this memory for more than one month. We use the sticky tape avoidance test to demonstrate that striatal LTP emerges in vivo after one-shot learning; the observed patterns of activity in vivo suggest that eCB-LTP mediates one-shot learning, which we corroborate both *ex vivo* and through computational modeling. Consistent with this hypothesis, conditional knock-out mice abolishing eCB-LTP show impaired one-shot learning. These results highlight the importance of non-classical plasticity mechanisms in supporting memory formation after a single brief experience.

---

One-shot learning, the ability to acquire a memory from a unique and brief exposure to a new experience, is crucial for behavioral adaptation. This widespread phenomenon is involved in multiple learning processes, eg. spatial learning, recognition memory, complex episodic-like associations, instrumental learning in response to threat or appetitive stimuli, and inference learning during new word acquisition. All have in common that a single and brief experience, occurring on a second or subsecond timescale, is sufficient to trigger a long-term memory^1^. Despite its behavioral significance, the exploration of the neuronal mechanisms of one-shot learning is still in its infancy as it faces two main challenges. The first comes from the scarcity of rodent behavioral assays of naturalistic one-shot learning, outside of spatial learning and fear-based tasks. The second difficulty comes from the small number of action potentials elicited during a one-shot experience^2–7^. Such a limited neuronal activity typically contrasts with the Hebbian paradigm that postulates that multiple repetitions of coincident neural activity are required for stable plasticity expression, as what is likely happening during incremental trial-and-error learning. Interestingly, some plasticity rules relying on sparse activity or single burst firing have been uncovered *in vitro*^8–13^ and *in vivo*^14^. At this point, only the hippocampal behavioral timescale synaptic plasticity relates to behaviorally-relevant phenomena, i.e. the formation and remapping of place fields during spatial navigation^14,15^. We are thus critically lacking knowledge on the plasticity rules and circuits underlying other forms of one-shot learning.

The dorsal striatum, a major circuit involved in sensorimotor and instrumental learning which can manifest in one-shot, has not yet been investigated through the lens of one-shot learning. Indeed, several studies have emphasized the critical involvement of the dorsolateral striatum (DLS) in late learning stages^16–18^. While it has been hypothesized that one-shot vs. incremental learning should be implemented in separate circuits^19^, long-lasting modulation of neuronal activity has also been reported in the DLS in early learning stages^20–24^, or following rapid adaptation to abrupt changes in task contingencies^25,26^. While these behavioral paradigms are not one-shot learning tasks *per se*, they nevertheless tend to indicate that striatal projecting neurons (SPNs) can rapidly form novel associations from cortical inputs, and participate in newly adapted motor programs^27,28^. In addition, our previous work has shown that at cortico-striatal synapses, few spikes evoke an endocannabinoid-mediated long-term potentiation (eCB-LTP) while more intense and repetitive stimulation induces an NMDAR-LTP *in vitro* ^12,29–32^. These findings suggest that the same circuit – DLS – could mediate both one-shot and incremental learning undergoing synaptic plasticity dependent on distinct molecular signaling pathways.

Our results establish that a cortico-striatal LTP emerges *in vivo* upon a spontaneous one-shot learning experience in a novel behavioral test in mice, only if brief (less than 20 seconds). We find that the endocannabinoid system is a key molecular pathway of one-shot learning. Indeed, our data analysis based on a mathematical model of the synapse shows that the cortico-striatal network displays behaviorally-correlated activity patterns consistent with eCB-LTP induction. By examining the causal implication of the eCB-LTP in one-shot learning, we observe that conditional knock-out mice abolishing eCB-LTP expression show impaired one-shot learning. Together our data demonstrate that one-shot learning in the dorsal striatum is mediated by an non-classical plasticity mechanism, dependent on the endocannabinoid system.

## Novel one-shot learning test induces robust memory in rodents

To characterize the engram at play during one-shot learning in mice, we set up a novel behavioral test, the sticky tape avoidance (STA) test (**Fig. 1**). STA relies on spontaneous and unconstrained behavior, involving no reward nor penalty. At *familiarization*, the mouse explores an arena in which a piece of loose sticky tape has been positioned on the floor near the arena walls, with the sticky face up. The mouse rapidly sticks onto the tape, carries it away and then removes it. In the next *retrieval* session, occurring at least 24 hours after familiarization, the mouse is placed again in the arena with a new piece of sticky tape positioned at a different location. As a control, the same test was performed with a piece of non-sticky tape replacing the sticky tape (n-STA test). Comparing mice behaviors between familiarization and retrieval allows assessing learning performance: the manifestation of avoidance behavior in the vicinity of the sticky tape at reteival reflects the formation of a long-term memory subsequent to a single, typically brief, exposure to a sticky tape, namely a successful one-shot learning (**Fig. 1a**). Avoidance behavior, clearly visible in the trajectories of the mice in the arena (**Fig. 1a**), can be quantified through the significant decay of the mouse occupation probability and average speed in the vicinity of the sticky tape (n=56 mice), in sharp contrast with the n-STA control experiment where neither the occupation probability nor speed was affected around the location of the non-sticky tape (n=22 mice) (**Fig. 1b** and **Extended Data Fig. 1a**).

**Figure 1:**
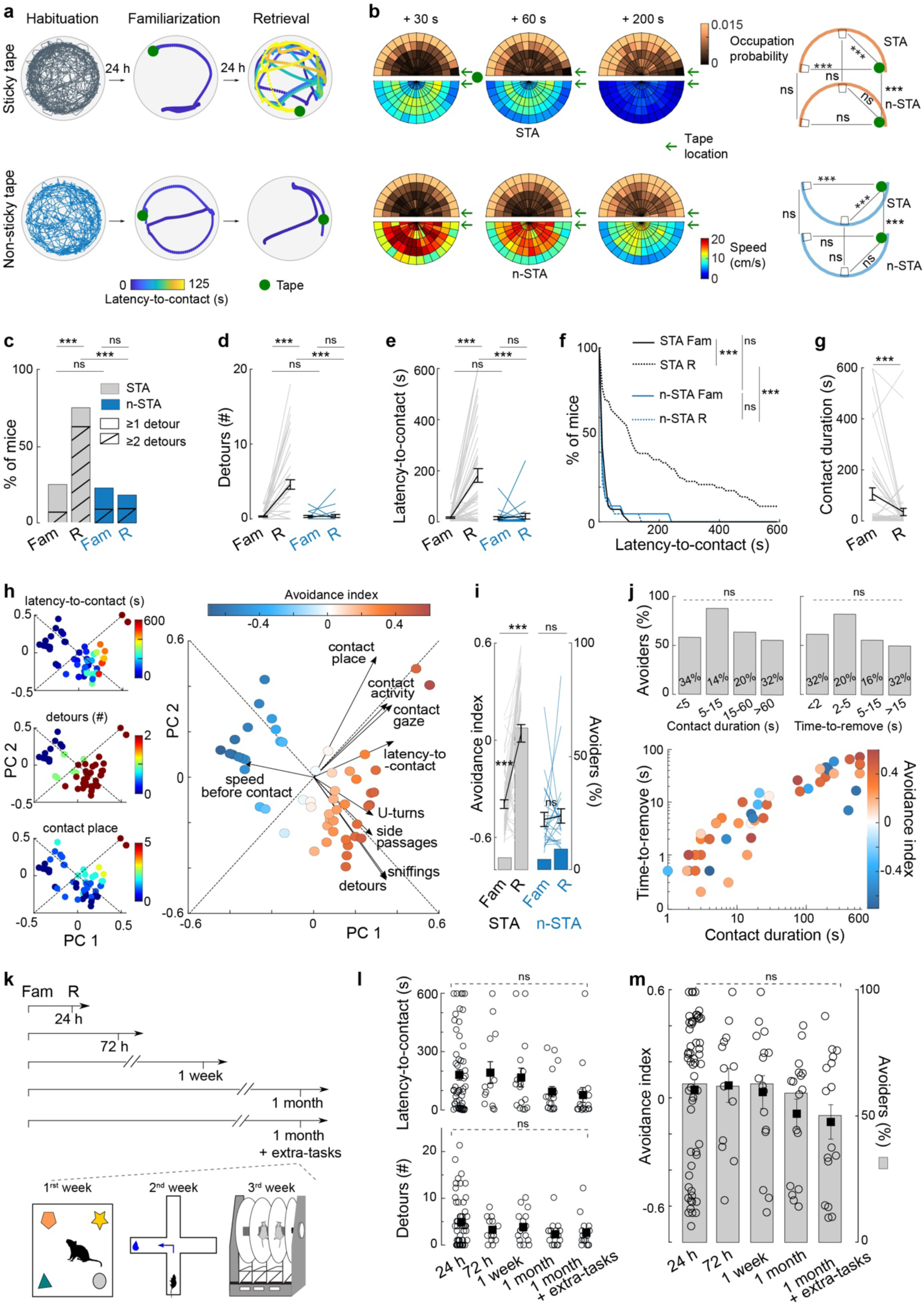
Induction of one-shot learning with the sticky-tape avoidance test. **a**, Individual mouse trajectory in STA and n-STA tests. **b**, Average mouse trajectory in n-STA and STA during retrieval. 2D cumulative heat-maps of the mouse occupation probability and average speed. For STA, only mice with latency-to-contact superior to each timepoint are included (n=36, 34, 19); n-STA: n=22. Green arrow: tape position. Trajectories are symmetrized relative to the tape position. At retrieval, mouse occupation probability and speed are affected by the sticky tape, indicative of one-shot learning, but not in n-STA. Linear Mixed Model with Tukey post hoc tests. **c-g**, Mice subjected to STA (n=56) showed increased numbers of detours (**c**,**d**) and latency-to-contact (**e**,**f**) with decreased contact duration (**g**) between familiarization (Fam) and retrieval (R), while n-STA mice (n=22) show no changes. McNemar test (c), repeated-measures ANOVA with Tukey post hoc test (d,e), log-rank test (f) and paired two-sample t-test (g). **h**, Individual mice scoring quantified from a PCA-based avoidance index, computed from a linear combination of the first two principal components (PCs), overlaid with parameter values (left) or retrieval avoidance index (middle). Vectors quantify the contribution of each parameter to the first two PCs. **i**, Increase in avoidance index between familiarization and retrieval, and in the percentage of avoiders (bars). Paired two-sample t-test and McNemar test. **j**, Brief contact duration and time-to-remove (<5 and <2 seconds, respectively) at familiarization are sufficient for one-shot learning, with avoidance index similar to longer contacts. Avoiders for different time bins of contact duration and time-to-remove (top); Percentage: proportion of mice belonging to each time bin. Avoidance index as a function of contact duration and time-to-remove (bottom). Paired two-sample t-test and Chi-squared test. **k,l,m**, Robustness over time of one-shot learning, assessed with (**k**) four increasing time retention intervals (24 hours, n=56 mice; 72 hours, n=13; 1 week, n=16; 1 month, n=17), and over time (1 month) with extra-tasks (n=16 mice) (novel object recognition, crossmaze and rotarod). Latency-to-contact and detours (**l**), avoidance index and avoiders (bars) (**m**), do not show difference across these 5 conditions. One-way ANOVA (m,n) and Chi-squared test (n). Statistics are provided in the *Methods*. \**p<*0.05; \*\**p<*0.01; \*\*\**p<*0.001.

At familiarization, the tape constituted a novel object that was quickly explored and contacted by most of the mice, with no or very few (one or two) detours, regardless of the nature of the tape, sticky or non-sticky indicative of the non-repelling nature of the sticky tape (**Fig. 1c-f, Extended Data Fig. 1b,c**). At STA retrieval, 24 hours after familiarization, a majority of mice exhibited an avoidance behavior, with on average a major increased in number of detours and an elevenfold increase in the latency-to-contact (**Fig. 1c-f** and **Extended Data Fig. 1d**). In detail, 63% of mice exhibited at least two detours, compared to 7% at familiarization (McNemar test, *p<*0.001). Males and females displayed a similar behaviors (**Extended Data Fig. 1e**). In addition, mice exhibited a better efficiency at sticky tape removal, as measured by a faster discarding of the tape at retrieval (**Fig. 1g**). Conversely, in n-STA, the number of detours and latency-to-contact were not significantly different between familiarization and retrieval (**Fig. 1c-f**). Altogether, the increase in both the number of detours and the latency-to-contact, specific to the STA test, indicates that long-term one-shot memory can be formed out of a one-shot encounter with a sticky tape and validates our novel behavioral paradigm as a one-shot learning test for mice.

To more precisely score individual mouse avoidance at retrieval, we performed an in-depth analysis of the mouse behavior which captures its non-trivial multifarious nature (*Methods, Supplementary Information*). To this purpose, we designed a PCA-based avoidance index taking into consideration nine parameters characterizing periods before contact with the sticky tape (latency-to-contact, detours, sniffings, U-turns, side passages) and at contact (speed in the 2 seconds preceding contact, activity, gaze and body part contacting the tape) (**Fig. 1h**, **Extended Data Fig. 2a,b,c,** *Methods, Supplementary Information*). Recapitulating the above observations, the avoidance index sharply distinguished behaviors at familiarization and retrieval (-0.39±0.03 *vs*. 0.04±0.06, paired t-test, *p<*0.001)), yielding 63% of avoiders after a single contact with the tape (**Fig. 1i**).

One-shot learning is defined not only by the singleness of the experience but also its brevity. To explore the lower temporal limits of one-shot learning in the STA test, we investigated whether the avoidance index at retrieval correlated with the duration of contact with the sticky tape and the time-to-remove (*i.e.* the cumulative duration of active tape removal by mice) at familiarization. We found that very brief contacts, below 5 seconds, induced avoidance in 58% mice, similarly to other ranges of contact duration (**Fig. 1j**). Surprisingly, the longest contact range (from 60 to 600 seconds) did not induce a higher percentage of avoiders (56%) compared to brief contacts (**Fig. 1j**), showing that mice need only a few of seconds in contact with the sticky tape to learn STA. Hence, the STA test enables to examine core features of one-shot learning, typically observed in the wild but rarely captured in existing behavioral paradigms.

To assess the robustness over time of this one-shot learning, three groups of mice were subjected to increasing time intervals between familiarization and retrieval. We observed that after 72 hours, one week or even up to one month, mice still exhibited marked avoidance of the sticky tape with elevated latency-to-contact and detours, and an avoidance index range similar to the 24 hours delay (**Fig. 1k-m**). At the population level, we did not observe significant differences between the avoider rate after 24 hours delay (63%, n=56 mice) and longer delays (72h: 62%, n=13; 1 week: 63%, n=16; 1 month: 59%, n=17; χ^2^-test, *p=*0.928), regardless of contact duration (**Fig. 1k-m, Extended Data Fig. 2d**). To further test the robustness of one-shot learning, we subjected mice (n=16) to three striatal-dependent learning tasks between familiarization and the one-month delayed retrieval (i.e. accelerating rotarod, cross-maze and novel object recognition). Here again, no significant difference in avoidance index and avoider rate were observed relative to the one-month delay without interleaved tasks. Good and bad fast learners for STA (with high and low avoidance index, respectively) performed equally well on the three additional tasks (**Extended Data Fig. 2e**). These results demonstrate that STA-induced one-shot memory is robust over time and to interfering long-term memory tasks. Overall, STA constitutes a novel one-shot learning paradigm in mice, which relies on the spontaneous behavior of mice, captures quintessential manifestations of one-shot learning and induces lasting and robust long-term memories.

## One-shot learning induces *in vivo* cortico-striatal LTP

To explore the neuronal substrate of one-shot learning, we monitored cortico-striatal plasticity during STA. Indeed, STA relies on learning to select an appropriate motor behavior in response to a sensory stimulus, a function supported by cortico-striatal circuits^33,34^. To test whether cortico-striatal plasticity was induced after a one-shot experience with the sticky tape, we monitored possible changes in the amplitude of DLS responses to cortical stimulations. To this end, we opto-stimulated somatosensory cortical terminals expressing channelrhodopsin (ChR2) and recorded the evoked local field potential (ChR2-LFP) in DLS with a chronically-implanted optrode (**Fig. 2, Extended Data Fig. 2f, Extended Data Fig. 3a**). We confirmed that the late-phase of ChR2-LFP reflected synaptic currents, since it was reduced by inhibition of AMPA and NMDA receptors, indicating its excitatory nature, and fully abolished in the absence of spiking activity after blockade of voltage-dependent sodium channels (**Fig. 2, Extended Data Fig. 3b, c**). Hence, the amplitude of the late phase of ChR2-LFP (normalized by early phase amplitude) was used as a proxy for cortico-striatal plasticity *in vivo*^35^ (*Methods*).

**Figure 2:**
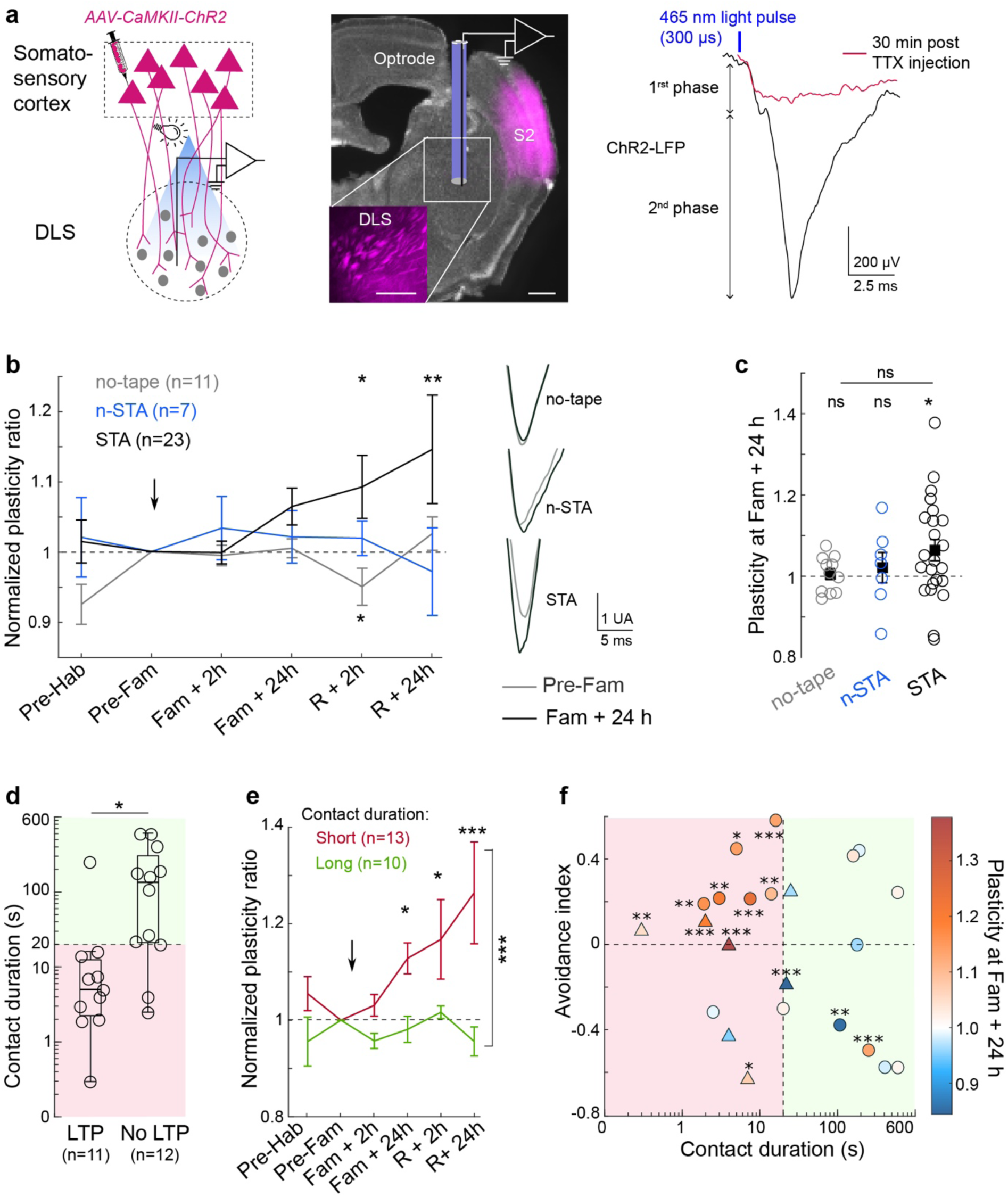
A single and brief exposure to the sticky tape induces *in vivo* cortical-striatal plasticity. **a**, ChR2-LFP monitoring to assess *in vivo* cortico-striatal plasticity (*Methods*). Virus expression (magenta) and optrode, zooming on somatosensory cortical terminals in the dorsal striatum. Scale bars: 0.5 mm. Illustrative ChR2-LFP raw trace from a single light pulse stimulation. ChR2-LFP displayed an early phase (latency-to-peak: 0.77±0.04 ms, n=41), due to photoelectric artifact and light-evoked ChR2 currents, and a late phase (latency-to-peak: 2.7±0.1 ms) accounting for synaptic currents. *In vivo* injection of TTX (1µM, 1µL) shows an abolishment of ChR2-LFP. **b**, Averaged time courses of plasticity ratio (normalized by the ChR2-LFP amplitude at pre-familiarization) show specific LTP from 24 h after sticky tape contact (STA), but not for the no-tape and n-STA conditions. Representative ChR2-LFP traces at pre-familiarization and familiarization +24 h. Linear Mixed Model with Tukey post hoc test. **c**, Scatter plot of averaged plasticity ratio at familiarization + 24 h for individual mice. One-way ANOVA and one-sample t-test. **d**, Contact duration with the sticky tape at familiarization discriminates between mice exhibiting or not ChR2-LFP potentiation (dashed line: threshold value of 20 seconds; pink and green shades indicate short and long contact duration, respectively). Two-sampled t-test. **e**, Averaged time courses of plasticity ratio for STA mice depending on contact duration (threshold ≤20 seconds). Linear Mixed Model with Tukey post hoc test. **f**, Plasticity ratio at familiarization +24 h (color-coded) as a function of STA avoidance index and contact duration for individual mice shows that all mice displayed ChR2-LFP potentiation after short contact duration associated with high avoidance index. Circles and triangles represent mice after 24 hours or 48 hours/1-week delays, respectively, between familiarization and retrieval. One-sample t-test and ANCOVA. Statistics are provided in the *Methods*. \**p<*0.05; \*\**p<*0.01; \*\*\**p<*0.001.

Chronic *in vivo* recordings of cortico-striatal plasticity were performed in three groups of mice, either in an empty arena (no-tape, n=11) or subjected to the n-STA (n=7) or STA (n=23) tests (**Fig. 2b**,**c****, Extended Data Fig. 3d**). In both control groups (no-tape and n-STA), we did not observe any change of ChR2-LFP time course. In contrast, in STA, a long-term potentiation (LTP) of ChR2-LFP was expressed 24 hours after familiarization and persisted at least 24 hours after retrieval (one-sample t-test, *p=*0.023) (**Fig. 2b**,**c****, Extended Data Fig. 4a**). This LTP emerging at cortico-striatal synapses after familiarization and before retrieval likely participates to the engram of one-shot learning. Surprisingly, only half of the mice displayed LTP 24 hours after the first contact. To examine the origins of this dichotomy, we correlated *in vivo* plasticity to behavior at familiarization and retrieval (**Extended Data Fig. 4b**). We observed that 91% mice expressing LTP (n=11) had a short contact duration with the sticky tape (2 to 20 seconds), whereas 75% mice showing no LTP (n=12) displayed a long contact (>20 seconds) (independent t-test at +24 hours, *p=*0.03; Linear Mixed Model, *p=*0.003) (**Fig. 2d**,**e****, Extended Data Fig. 4b-f**). LTP was found for time-to-remove ranging from 0.2 to 15 seconds (**Extended Data Fig. 4c**). Our results hence demonstrate the emergence of LTP in cortico-striatal circuits subsequent to a very brief experience. Moreover, this LTP appears specific to brief experiences, as mice exhibiting a long contact duration (>20 seconds) at familiarization, failed to express this LTP, even when testing 24 hours after a second exposure (*i.e.* retrieval +24 hours) (**Extended Data Fig. 4g**). Finally, all mice that displayed short contact duration and LTP showed a high avoidance index (ANCOVA, *p=*0.041) (**Fig. 2f, Extended Data Fig. 4h**). For long contact duration (>20 seconds), high behavioral avoidance could be observed despite the lack of LTP in DLS, suggesting that this phenomenon is specific to one-shot learning, and that longer exposures involve other brain circuits.

Overall, cortico-striatal LTP in DLS is elicited by a brief, single and salient experience. A very brief exposure, as low as 2 seconds, is sufficient to trigger synaptic plasticity, which correlates with one-shot learning at the behavioral level (**Fig. 2f**).

## Endocannabinoids mediate LTP induction in one-shot learning

According to the above observations that LTP is elicited exclusively during short contacts and not upon longer contacts, we hypothesize that the signaling pathways underlying the cortico-striatal LTP induced by one-shot learning should be activated with relatively few spikes or bursting events, and be switched off upon longer and repetitive stimulation. Such plasticity is evocative of a particular type of LTP induced by a spike-timing-dependent plasticity (STDP) paradigm^36^ mediated by endocannabinoids (eCB-LTP) that we identified *in vitro* in DLS^12,29,30^. Indeed, eCB-LTP was induced by 10 to 15 STDP pairings (paired activity on either side of the cortico-striatal synapse, in a post-pre-synaptic temporal sequence) and was no longer expressed after 50 pairings. This eCB-LTP requires the activation of presynaptic cannabinoid type-1 (CB1R) and dopamine type-2 (D2R) receptors^30^. Here, we confirmed that eCB-LTP could be elicited in DLS with only 15 pairings using patch-clamp recordings of individual SPN in mouse brain slices (*Methods*). We further found that no LTP could be elicited in the dorsomedial part of the striatum (DMS) following the same plasticity induction protocol, and even for higher pairing frequency (2.5Hz) (**Extended Data Fig. 5a**). These results indicate that cortico-striatal synapses in DLS may be more prone than DMS to plasticity when few spikes occur.

To test whether the spiking activity patterns in cortico-DLS circuits occurring during STA could be compatible with eCB-LTP induction, we performed *in vivo* Neuropixel 1.0 recordings of somatosensory cortical and DLS neurons, and used these spike trains as an input to stimulate a detailed mathematical model of the cortico-striatal synapse (**Fig. 3**). Neuronal recordings were performed in head-fixed mice navigating inside the air-lifted Mobile HomeCage (**Fig. 3a**, *Methods*). In a subset of mice (n=4 out of 9), contact durations at familiarization were short, from 2.5 to 8 seconds, and were accompanied by detours during the retrieval session, 24 hours later (**Fig 3a**, **Extended Data Fig. 5b**). To estimate the level of neuronal activity during contact with the sticky tape, we quantified the firing rates and absolute number of spikes of cortical and striatal neurons. They remained low for the majority of neurons across the four mice (median ranging from 1 to 14 spikes in the cortex and from 1 to 35 in DLS). (**Fig. 3b**). Such a sparse firing activity of cortical and DLS neurons is therefore compatible with the specifics of eCB-LTP

**Figure 3:**
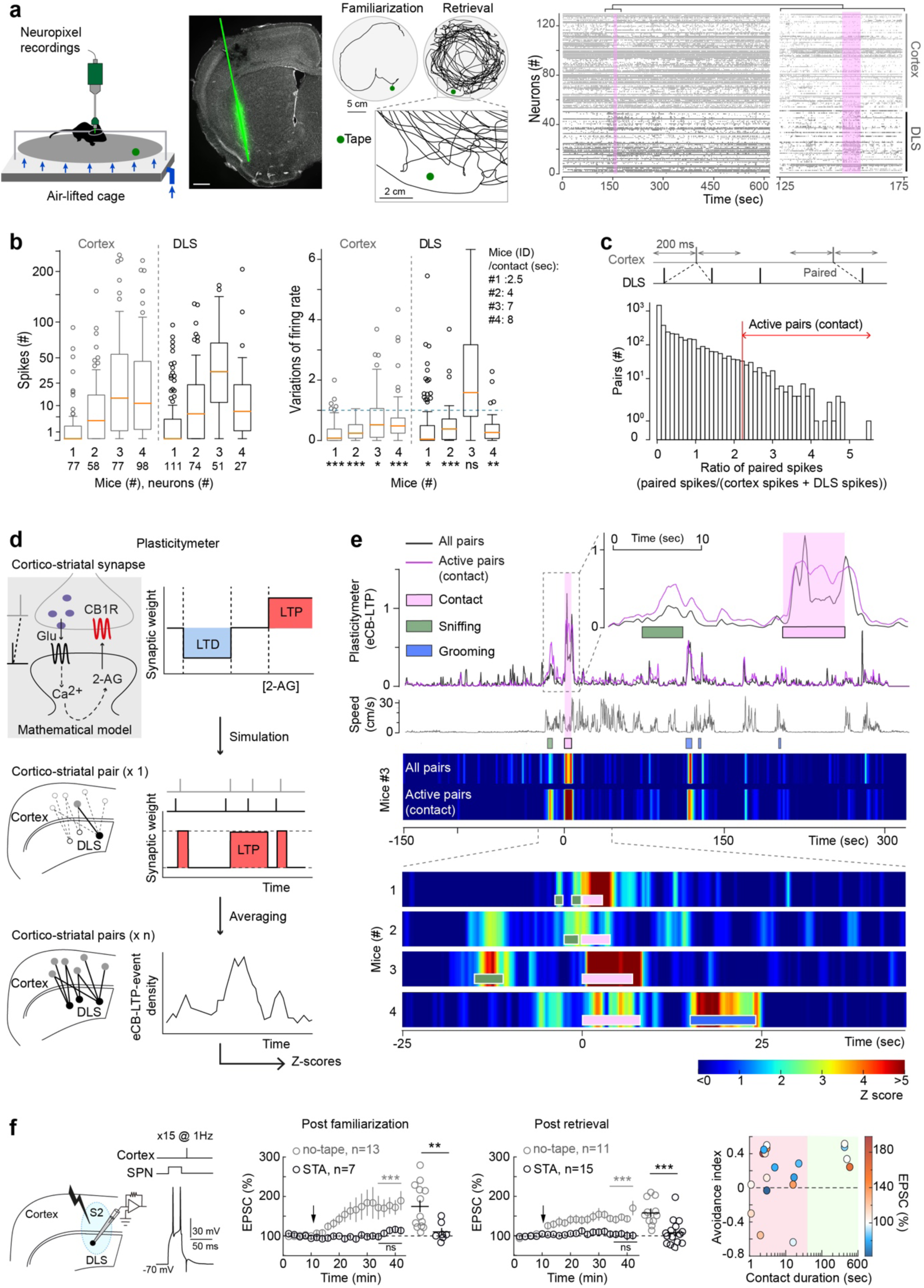
Cortico-striatal activity induced upon a brief contact with the sticky tape engages endocannabinoid-mediated LTP. **a,** Acute Neuropixel 1.0 recordings in head-fixed mice in the MobileHomeCage (*Methods*). Coronal brain slice showing the Neuropixel trajectory, targeting somatosensory cortex and DLS. Scale bar: 0.5mm. Illustrative mouse trajectory at familiarization and retrieval with insert-zoom on the detours close to the tape at retrieval. Example (mouse#3) raster plot displays the spiking activity of cortical (grey) and DLS (black) units obtained after spike sorting (Kilosort-2.5) and manual curation (Phy-2.0); Contact period: overshadowed in pink. **b,** Boxplots of the absolute number of spikes emitted by cortical and DLS units, during short contacts (from 2.5 to 8 seconds), indicate a sparse activity (average median number of spikes in cortex: 7.25 and DLS: 12). Boxplots of the ratio of the firing rates during contact *vs*. baseline show that the contact does not induce an increase in number of spikes compared to the 95^th^ centile observed during baseline. **c**, Identification of pairs of cortical and striatal neurons showing a high number of correlated spikes (±200ms) during tape contact. *Active pairs* correspond to the 5% of pairs showing the highest number of correlated spikes. **d,** The *plasticitymeter* uses a mathematical model of endocannabinoid-mediated STDP driven by paired spike trains from individual cortical and striatal units to predict the occurrence of eCB-LTP induction events. Simulations can be performed across all combinations of cortico-striatal pairs of neurons or across the active pairs. **e,** Plasticitymeter for a representative mouse (#3) and behavioral parameters, including mouse instantaneous speed, interactions with the tape or following contact. Plasticitymeter z-scores across all pairs or only active pairs show that the contact with the sticky tape is associated with a significant increase in eCB-LTP induction events (z-score >2 for every animal at contact time). Z-score maps are aligned to the start of contact. **f,** *Ex vivo* patch-clamp occlusion experiments in DLS SPNs of mice subjected to STA or no-tape conditions. STDP protocol: 15 cortico-striatal post-pre pairings at 1Hz (Δt_STDP_∼-15ms). The endocannabinoid system in DLS SPNs is engaged after contact with the sticky tape but not in the no-tape condition, after familiarization and retrieval (right). One-sample and two-sample Wilcoxon tests. E*x vivo* plasticity related to avoidance index at retrieval (R) and contact duration at familiarization (Fam). Contact duration (<20 seconds: pink and >20 in green). Statistics are provided in the *Methods*. \**p<*0.05; \*\**p<*0.01; \*\*\**p<*0.001.

To examine more precisely whether the firing patterns of cortical and striatal units could elicit eCB-LTP, we developed a mathematical model of *in vivo* cortico-striatal STDP building upon a previous model version validated for mouse brain slice recordings^12,29–31,37^. This mathematical model expresses the kinetics of the main molecular species involved in the signaling pathways of eCB-dependent LTP and LTD as a function of presynaptic and postsynaptic spikes (*Methods*). In the model, expression of endocannabinoid plasticity depend on CB1R activation level: eCB-LTD for intermediate endocannabinoid levels and eCB-LTP for large endocannabinoid levels. We used the *in vivo* single-unit cortical and striatal spike trains for each mouse as inputs to the model and estimated resulting CB1R activation and eCB-LTP induction events, referred to as “plasticitymeter” (**Fig. 3c-e**, **Extended Data Fig. 5c**; *Methods*). Active pairs – exhibiting the largest relative increase in correlated activity during contact (**Fig. 3c**; *Methods*) were associated with high plasticitymeter levels, particularly during behaviorally-relevant states: contact with the sticky tape, active locomotion, grooming and sniffing of the sticky tape (**Extended Data Fig. 5c,d, Fig. 3d-e)**. In particular, most eCB-LTP events were concentrated around the contact with the sticky tape in all mice, associated with elevated z-scores (> 2 for all mice in at least 10% of the first 10 seconds after contact onset; also at least three times larger than during baseline). This observation remained consistent when selecting cortico-striatal neuron pairs with a smaller increase in their correlated activity during contact, and even in some mice when taking into account all possible pairs (**Extended Data Fig. 5e).** Overall, the plasticitymeter model shows that the firing patterns of cortical and striatal neurons are compatible with the induction of eCB-LTP during short contacts with the sticky tape.

We thus set out to test experimentally whether eCB-LTP was engaged right after STA. For this purpose, we performed *ex vivo* patch-clamp occlusion experiments in acute brain slices and tested eCB-LTP induction in DLS 30 min after familiarization or retrieval in STA or no-tape conditions (**Fig. 3f**). After retrieval, 70 % of STA mice exhibited a behavioral avoidance (n=15 mice with a similar % of avoiders than in **Fig. 1**). Among mice showing avoidance, no plasticity could be elicited in SPNs after applying eCB-LTP induction protocol (n=15 SPNs), in sharp contrast to the no-tape condition in which potent LTP was evoked (n=11 SPNs). To examine the plasticities induced after a one-shot experience, we applied the same protocol immediately after familiarization; we found that LTP was occluded in STA (n=7 SPNs), but not in the no-tape condition (n=13 SPNs). LTP occlusion was observed for a wide range of contact duration at familiarization, including very short ones down to 2 seconds (**Fig. 3f**). Our occlusion experiments strengthen the hypothesis that the endocannabinoid system is engaged right after a one-shot experience.

To conclude, *in vivo* recordings of cortical and striatal neurons showed low levels of activity during STA, compatible with the induction of eCB-LTP as suggested by mathematical modeling. Since eCB-LTP is also occluded *ex vivo* after STA, these data suggest altogether that endocannabinoids are recruited during one-shot learning and are the substrate of the *in vivo* LTP observed in **Fig. 2**.

## Impaired one-shot learning in eCB-LTP knock-out mice

To investigate whether endocannabinoids are required for one-shot learning, we took advantage of the fact that activation of CB1R or D2R specifically at presynaptic cortical afferents is required for eCB-LTP expression^30^. We therefore generated conditional eCB-LTP knock-out, by virally expressing Cre recombinase in the somatosensory cortex of the transgenic CB1R^flox/flox^ or Drd2^LoxP/LoxP^ mice (CB1R-Cre or Drd2-Cre mice) (**Fig. 4a-j**); mice expressing GFP instead of Cre served as control animals (CB1R-control or Drd2-control mice). We verified with whole-cell recordings that eCB-LTP was impaired *in vitro* in CB1R-Cre or Drd2-Cre mice (**Fig. 4b,g**). To do so, we applied 15 cortico-striatal STDP pairings in brain slices from naive mice and observed an absence of LTP in SPNs from CB1R-Cre and Drd2-Cre mice (n=10 and 8 SPNs, respectively), whereas LTP was induced in both CB1R-control or Drd2-control mice (n=9 and 9 SPNs, respectively).

**Figure 4:**
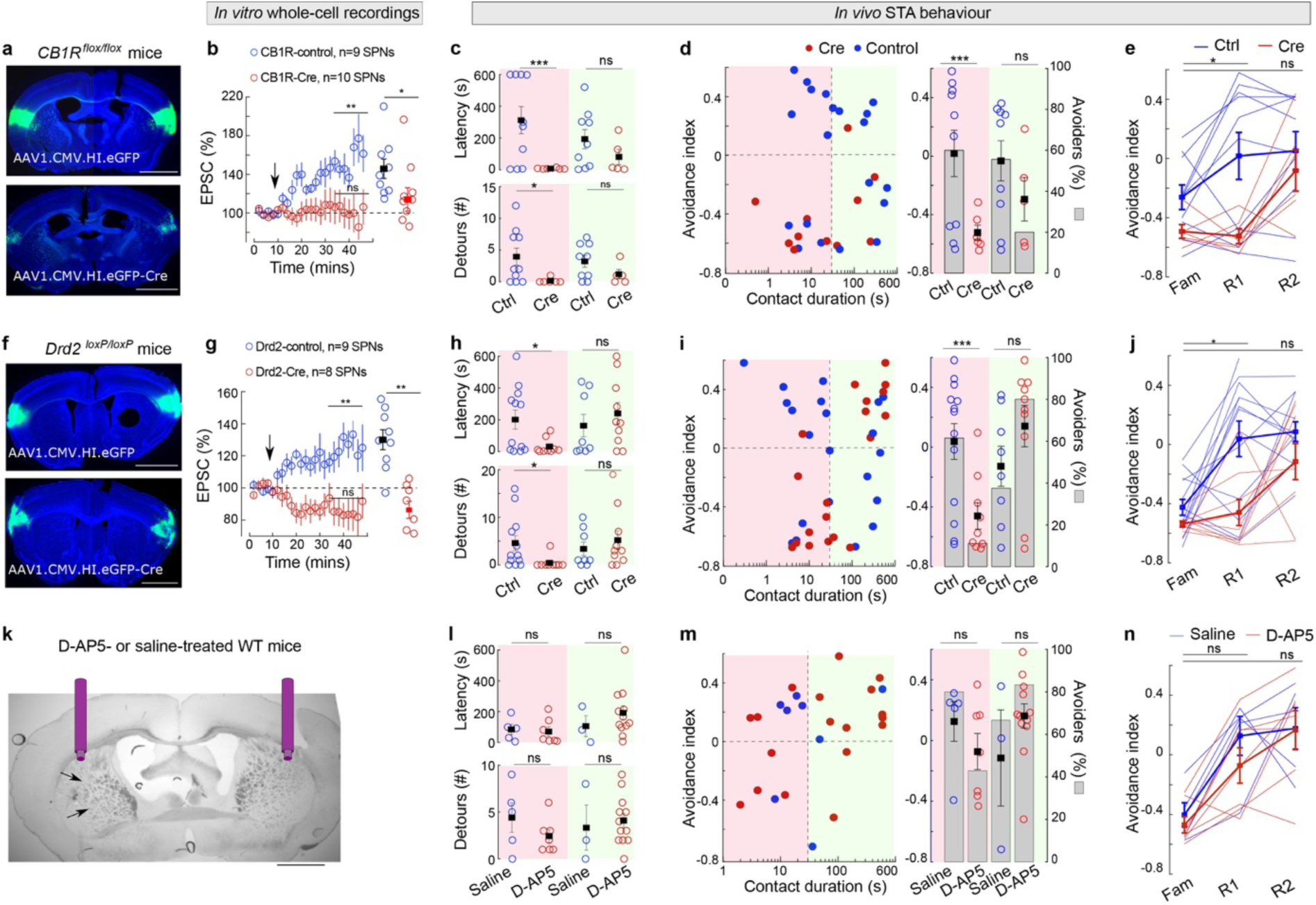
One-shot learning is impaired in cortico-striatal presynaptic CB1R or D2R knock-out mice but not under NMDAR inhibition. **a**-**j**, Conditional CB1R^flox/flox^ and Drd2^LoxP/LoxP^ mice show impaired STA one-shot learning. **a**,**f**, Bilateral virus expression (AAV1.CMV.HI.eGFP for control or AAV1.CMV.HI.eGFP-Cre for cre mice) targeting somatosensory S2 cortex. Scale bars, 0.5 mm. **b**,**g**, *Ex vivo* whole-cell recordings on naïve conditional knock-out mice show impaired eCB-LTP expression after STDP 15 post-pre cortico-striatal pairings (1Hz, Δt_STDP_∼-15ms). Averaged time-courses of STDP plasticity for control and Cre mice. Scatter plots: changes in EPSC amplitude 30 min after STDP pairings. One-sample and two-sample Wilcoxon test. **c**,**h**, Latency-to-contact and detours. Two-sample Welch-test (for short contact duration, ≤30 seconds, because of unequal variance) and t-test (long contact duration >30 seconds). **d**,**i**, Avoidance index (circles) and avoiders (bars) are low in Cre mice for short contact duration, but remain high for long contact duration. Two-sample Welch and t-test, Chi-squared test. **e**,**j**, Time courses of avoidance index at familiarization, retrieval-1 and -2 sessions for short contact duration mice. Linear Mixed Models. **k**-**n**, Wild-type mice injected bilaterally with D-AP5 (500 µM) or saline 45 minutes before STA familiarization. **k**, Cannulas implanted in DLS and injection sites (arrow). Scale bar, 0.25mm. Latency-to-contact and detours (**l**), avoidance index (circles) and avoiders (bars) (**m**), show no change for short and long contact duration in D-AP5- and saline-injected mice. Two-sample Welch and t-test, Chi-squared test. **n**, D-AP5- and saline-injected mice, with a short contact duration at familiarization, display similar time course of the avoidance index at familiarization, and first and second retrieval (retrieval-1 and retrieval-2). Linear Mixed Model. Statistics are provided in the *Methods*. \**p<*0.05; \*\**p<*0.01; \*\*\**p<*0.001.

Based on the above observations, we hypothesized that one-shot learning should be impaired in eCB-LTP knock-out mice. To test our hypothesis, we examined the behavior and learning performance of CB1R and D2R conditional knockout during STA (n=11 CB1R-Cre and n=19 CB1R-control mice; n=19 Drd2-Cre and n=21 Drd2-control mice). Overall, at familiarization, all control and Cre mice showed similar behavior (**Extended data Fig. 6**). At retrieval, regarding CB1R^flox/flox^ mice, we observed reduced latency-to-contact and number of detours for short contact duration in CB1R-Cre, in contrast to CB1R-control mice (independent t-tests, latency-to-contact: *p*=0.006; detours: *p*=0.022; n=6 *vs*. 10) (**Fig. 4c**). Importantly, we found that none of the short contact duration CB1R-Cre mice avoided the tape (0% avoiders), in contrast to CB1R-control mice that performed similarly to wild-type mice with 60% of avoiders (avoidance index: independent t-test, *p=*0.008; avoider rate: χ^2^-test, *p=*0.016) (**Fig. 4d**). Similarly, short contact duration Drd2-Cre mice showed reduced latency-to-contact and number of detours at retrieval, in contrast to Drd2-control mice (independent t-tests, latency-to-contact: *p=*0.015; detours: *p=*0.027; n=9 *vs.* 13) (**Fig. 4h**). In addition, avoidance index and avoider rate were very low in short contact duration Drd2-Cre mice, compared to Drd2-control (avoidance index: independent t-test, *p=*0.006; avoider rate: χ^2^-test, *p=*0.018) (**Fig. 4i**). On the opposite, long-contact duration Drd2-Cre (n=10) exhibited a clear avoidance behavior, with similar performance as Drd2-control mice (n=8). Overall, these data show that one-shot learning is prevented in animals experiencing a short contact duration with the sticky tape when eCB-LTP is impaired, which validates our hypothesis. In addition, when we subjected mice to two subsequent retrieval sessions (with 24 hours interval), we found that if the first retrieval allowed to discriminate between knock-out and control mice for short contact duration, there was no longer a difference in avoidance index at the second retrieval (**Fig. 4e** and **4j**). This observation indicates that the endocannabinoid system at cortico-striatal synapses in the DLS is critical for one-shot learning, but not for forming memories after a second iteration of the same experience.

While we found that endocannabinoids can mediate an unconventional form of LTP at cortico-striatal synapses, the classical NMDA receptors have also been implicated in the induction of LTP *in vitro* and in DLS-dependent learning *in vivo*^21,38,39^. Therefore, to examine the contribution of NMDAR-mediated signaling in one-shot learning, we pharmacologically inhibited NMDA receptors in DLS 45 minutes before familiarization using bilateral D-AP5 micro-infusions (500μM); saline micro-infusions via cannulas 45 minutes before the familiarization served as control conditions (**Fig. 4k, Extended Data Fig. 7a**). At familiarization, D-AP5- and saline-injected mice exhibited a similar behavior (**Extended Data Fig. 7b**). At the first retrieval, we did not observe significant differences in D-AP5- and saline-injected mice (n=7 and 5 for short contact duration) for latency, number of detours, avoidance index and avoider rate (**Fig. 4l-n**). There was also no change in avoidance index in both conditions at the second retrieval.

In conclusion, genetic and pharmacological approaches demonstrate that endocannabinoids, but not NMDARs, are required for one-shot learning.

## Discussion

Our results establish a novel behavioural paradigm for studying one-shot learning in mice. The STA test relies on spontaneous and unconstrained exploration and allows for a rich repertoire of behaviours to be expressed, both during the first encounter with the novel stimulus, and on the following retrieval session. While the sticky tape is uncomfortable, it does not trigger at retrieval fear-related behaviours but rather more careful and repeated investigations while avoiding contact. The STA test induces robust long-term memory and relies on object recognition, and learning to appropriately select avoidance motor programs in the vicinity of the sticky tape. It therefore enlarges the spectrum of one-shot learning behaviours studied in laboratory, restricted until now to spatial learning^40,41^, or fear-based behavioral protocols such as fear conditioning or inhibitory avoidance tasks^42,43^. Furthermore, while current one-shot learning tasks rely on hippocampal and amygdala circuits, the STA test opens up the possibility of studying the involvement of the basal ganglia in one-shot learning, a yet unchartered research topic.

Our data reveal that one-shot learning in the dorsal striatum is mediated by an atypical plasticity mechanism, dependent on the endocannabinoid system. Endocannabinoids have been implicated in learning and memory^44,45^, mostly because of their powerful depressing function on synaptic plasticity^46^, but an increasing body of evidence has broadened this view and showed that endocannabinoids can also increase neurotransmitter release at multiple synapses of the brain^32,46^. Our data reinforce the evidence for endocannabinoid functions in mediating bidirectional control of synaptic plasticity^12,29,30,32,47^ and associate the eCB-LTP to a fundamental learning process, *i.e.* one-shot learning. In line with our results, eCB-LTP induction was shown to be more robust to spike timing jitter during the plasticity induction protocol *in vitro* than its NMDAR counterpart^31^. Such robustness to jitter may be especially required for one-shot learning, as a decrease in spike timing variability between activated neurons only emerges at the late learning stages^24,25,48^. In addition, our pharmacological investigation revealed that NMDAR signaling was not necessary for one-shot learning in our behavioural assay. The critical involvement of endocannabinoids, and not of NMDARs, in one-shot learning, is also consistent with the absence of change in AMPA/NMDA ratio, evidence for a presynaptic plasticity locus in DLS, together with no learning deficits during the initial rotarod trials after NMDAR blockade^21,22,49^ as well as indicating impaired learning in the inhibitory avoidance task upon CB1R but not NMDAR inhibition after a single training session^50,51^.

We observed a rapid *ex vivo* eCB-LTP occlusion, 30 minutes after mice first experienced the sticky tape, which affected 71% and 73% of cells randomly selected in brain slices after familiarization and retrieval, respectively. The rapid time course of plasticity occlusion could reflect early changes in phosphorylation states or the activation of effectors such as cAMP. The widespread feature of the occlusion can result from elevated DLS activation in the very first trials of a novel task^20,23,52,53^ and the nature of the eCB-LTP induction itself, which requires only little coincident cortico-striatal activity^12,29,30^. Hence, during the contact with the sticky tape, a large population of neurons may be subjected to activity patterns compatible with eCB-LTP induction. In this sense, endocannabinoid-mediated plasticity can be thought of as a priming mechanism, evoked upon a salient and brief stimulation. Subsequent phenomena involved in the refinement and consolidation of cortico-striatal plasticity could take place at later stages, and participate in the selection and stabilization of a specific subset of cells, which would underlie the memory engram. Such a scenario would be compatible with a previous report exploring the evolution of striatal activity patterns across learning trials^23^. Here, a more precise examination of eCB-LTP engagement along the behavioural test and of the nature of the LTP observed *in vivo* after the first retrieval could provide insight into the consolidation mechanisms of one-shot learning.

Our work focused on the DLS. Although DLS is largely described as being gradually engaged in later learning phases corresponding to habit formation^20,23,54,55,56^, mounting evidence suggest that DLS can be engaged from early training phases^55,56,22,24^. DLS is also involved in fast behavioral timescales, as it encodes sub-second behavior syllables^57^ and integrates fast dopamine transients^58^. Furthermore, we did not observe eCB-LTP *in vitro* in another striatal area, the DMS, yet classically engaged in the early learning phases and rapid adaptations to novel action-outcome associations^24,27,21,59^. Such differential sensibility to sparse firing activation could add another layer of functional segregation between the two circuits, which exhibit distinct plasticity rules^24^, and highlight the critical role of DLS in mediating one-shot learning upon very brief experience.

Indeed, our data also suggest that different synaptic substrates may underlie one-shot learning upon brief *vs*. long experience, and one *vs*. two-shot learning. DLS would be specifically involved in one-shot learning, upon a brief experience. Indeed, we did not observe LTP *in vivo* in mice experiencing long contact duration with the sticky tape at familiarization, and Drd2^LoxP/LoxP^ knock-out mice showed similar learning performance to control mice for long contact duration. Upon a second re-exposure, mice in which eCB-LTP was abolished no longer exhibited impaired learning. Whether DMS or cortical circuits take over in these two cases remains to be investigated. Such rapid transfer across brain regions has been reported in the inhibitory avoidance task in rats, showing that while the first trial required hippocampal activation, the second trial relied on DLS^50^.

In conclusion, we establish a novel behavioural paradigm for studying key features of one-shot learning in rodents. Our results reveal a novel neuronal mechanism underlying one-shot learning: LTP is elicited at cortico-striatal synapses upon a spontaneous and brief one-shot learning experience and the DLS endocannabinoid system is a key molecular pathway of such learning. Because of the richness of the behavioural repertoire expressed in our one-shot learning test, our data also unravel temporal and activity-dependent boundaries of the eCB-LTP involvment in one-shot learning in the DLS. Furthermore, our results revisit the roles of the DLS, classically associated to incremental learning, and shed new light on the functions of the bidirectional nature of endocannabinoid-based plasticity. This bidirectional endocannabinoid-plasticity may offer keys to interpret the functions of the endocannabinoid system and how one-shot leaning is impacted by cannabinoids, including endocannabinoid-based medicinal drugs.

## Methods

### Animals

All experiments were performed in accordance with local animal welfare committee (CIRB) and EU guidelines (directive 2010/63/EU). P7-10 weeks wild-type C57BL6/J mice (Charles River, France), *Drd2^loxP/loxP^* mice (JAX stock#020631) and *CB1R^flox/flox^* (gift from Marsicano’s laboratory, Bordeaux, France) of both sexes were used, group-housed on a standard 12h light/dark cycle and with *ad libitum* food and water. Homozygous *Drd2^loxP/loxP^* and *CB1R^flox/flox^* were obtained from heterozygous breeding mice (back-crossed with C57BL6/J mice). For surgeries, mice were anesthetized intraperitoneally with a mix of ketamine hydrochloride (67mg/kg, Imalgène, Mérial, Lyon, France) and xylazine (13mg/kg, Rompun, Bayer, Puteaux, France); during and after surgery, animals were kept at 34°C.

### The sticky tape avoidance test (STA)

STA was performed in a Plexiglas round-shaped arena of 30cm diameter, in a sound-attenuated room with a controlled light intensity of 50lx (350lx in the arena). STA was composed of at least three sessions (10 min each): (1) habituation to the empty arena; (2) familiarization (24 hours after habituation) in which a piece of one-sided sticky tape is inserted in the arena, and the experiment, of variable duration, runs until the mouse contacts the tape so that the tape adheres to a body part of the animal (**Extended Data Fig. 1b**), and extend during the period when the animal attempts to remove the tape; (3) retrieval session to assess learning avoidance. A second retrieval session was performed in a subset of animals (**Fig. 4**). Between familiarization and retrieval, variable retention intervals (24, 48, 72 hours, 1 week and 1 month) were tested. The sticky tape consisted of four hand-cut squares (0.8x0.8cm) of green insulating PVC tape, covering a total of 2x2cm surface, and was placed at 3cm from the wall at different locations between familiarization and retrieval (at least a difference of 90° angle). The tape was placed near the arena walls, where the mouse preferentially travels, to maximize a rapid encounter with the tape and underline the avoidance behavior. Mice were introduced at the opposite side of the tape location. The sticky tape was removed 15-30 seconds after it was taken off by the mouse. Another group of mice encountered the same tape but with the non-sticky face upwards (nSTA) on familiarization and retrieval sessions. In a third group (no-tape), mice were placed in the empty arena for three consecutive sessions. Animals were videotaped (30Hz frame rate, resolution: 1024x876px) and videos were analyzed offline

### Analysis of STA

Behavioral features were extracted from mouse trajectories and activities. Animal center of mass was extracted using a custom-made MATLAB script (version 2020b, The Mathworks, Natick, MA, USA) based on contrast differentiation; snout position was obtained using DeepLabCut^60^. Trajectories were smoothed using a Savitzky-Golay filter. The extraction of the behavioral features described below was performed by a combined analysis based on MATLAB routines and observation (blinded for transgenic mice).

#### Period from the entrance in the arena to the contact with the tape

We measured entrance speed (averaged instantaneous speed during the first two seconds inside the arena) and latency to 1^st^ passage of the mouse close to the tape (distance tape-body center <8cm). As the PCA analysis indicated that high speed before contact was associated with low avoidance index, we verified that the entrance speed was not correlated with the latency-to-contact at retrieval (Pearson’s correlation coefficient: r=0.032, p=0.735).

Mouse activities close to the tape were classified as follows:

- Sniffing (with distance tape-snout <4cm).
- Detour: deviation of mouse trajectory (angle <120°, with distance tape-center of mass <12cm).
- U-turn is a specific form of detour: approach-retreat sequence (angle <30°) with a retreat following the same pathway as the approach sequence, at ≧15cm/s speed.
- Side passage: rearing near the tape or walking past the tape (<8cm), without directed gaze/sniffing and without apparent change in trajectory (angle >120°). Stepping over the tape (without sticking to it) was included as a side passage.

Note that at familiarization, mice exhibiting clear signs of avoidance (latency-to-contact >180 seconds and detours ≧2 or latency-to-contact >100 seconds and detours ≧3) were excluded (n=18 mice, **Extended Data Fig. 1a**).

#### Period around the sticking up to the removal of the sticky tape

We evaluated parameters relative to the sticking modalities and attempts at removing the tape:

- the immediate pre-sticking speed (cm/s): averaged instantaneous speed during the last two seconds preceding contact.
- the immediate pre-sticking activity and gaze, labeled as follows: {0: sniffing, 1: retreat post-sniffing, 2: walking, 3: rearing, 4: others (*eg*. grooming/staying nearby still), 5: none*} and {1: directed gaze, 2: last-second gaze (mouse looked upwards while walking and directed its gaze only once very close to the tape), 3: distracted (mouse did not look in direction of the tape at the contact), 5: none*}.
- latency-to-contact (seconds)
- body part stuck to the tape: {0: snout; 1: forepaw, 2: hindpaw, 3: tail, 5: none*}.
- contact duration (seconds): total amount of time with the sticky tape glued to the mouse
- time-to-remove (seconds): cumulative durations of voluntary movements for tape removal; tape removed passively: 0.

* The indices describing contact configuration for initial body part, activity and contact gaze were ordered from 0 to 5, describing a decreasing level of engagement relative to the tape; the value of 5 was attributed to the subset of mice, which at retrieval did not stick to the tape.

#### Period after removal of the sticky-tape

The majority of mice successfully removed the sticky tape within the 10 minutes (except for 6% of mice); A subset (n=18 out of 74 mice) showing neophobic-like behavior were excluded (**Extended Data Fig. 1c**) because it prevents one-shot learning evaluation.

Complementary parameters and analyses are detailed in the *Supplementary Information*.

#### Distribution of mouse trajectory in space and time

To analyze mouse locomotor behavior relative to the tape we rotated the trajectories such that the tape was positioned at 0° angle. The polar coordinates of each mouse (center of mass) were binned (6 bins for the radius, ranging from 0 to 15 cm and 17 bins for the angle, from 0 to 180°). To avoid biases due to the particular choice of direction of movement of the mice breaking artificially the symmetry of the system, we only considered the absolute value of the angle. Occupation probabilities were normalized by the radius to account for the non-uniform surface across 2D bins. Average speed was obtained by averaging the instantaneous speed for every 2D bin. Occupation probabilities and averaged speed were calculated for each animal on the trajectories cumulated up to +30, +60 and +200 seconds as long the mouse did not stick to the tape (n=36, n=34, n=20, respectively, with n_total_=56 mice). In nSTA, since the tape was not removed at retrieval, the trajectories of all mice (n_total_=22) were considered for all timepoints. A Linear Mixed Model was fitted to the occupation probabilities and speed (restricted maximum likelihood algorithm). The model included the following factors: Cluster (nSTA vs. STA), Zone (3 predefined zones: around the tape, at 90° angle and opposite, of equal surface), Time (3 time points defined above). The cluster variable was the mouse ID. We have: Occupation Probability (or Speed) ∼ 1 + Cluster + Timepoints + Zone + Cluster:Timepoints + Cluster:Zone + Timepoints:Zone + Cluster:Timepoints:Zone + (1|Mouse ID) Similarly, the proportion of time spent immobile (instantaneous speed <2cm/s), near the walls (>2/3 of arena radius) and in half of the arena opposite to or containing the tape were estimated at various time points.

#### Avoidance index

To establish an avoidance index for each mouse, we selected a set of parameters that can be estimated for all mice at familiarization and retrieval: {latency before contact, number of detours, number of sniffings, number of U-turns, number of side passages, immediate pre-sticking speed, immediate pre-sticking activity and gaze, first body part contacting the tape}. Other possibly relevant parameters (% time spent near the walls, % time spent in the close vicinity of the tape, % time spent immobile) were excluded because they were highly correlated with the latency-to-contact and mostly meaningless for short latency-to-contact (**Extended Data Fig. 1a**). We clipped at 2 the number of sniffings, detours and side passages, and at 1 for U-turns. These thresholds were chosen to reflect qualitative differences in behaviors and to avoid biases associated with the latency to contact; the thresholds chosen maximized the difference in the proportion of mice between familiarization and retrieval for each activity (**Extended Data Fig. 1d**). A PCA was run on the z-scored parameters obtained at retrieval (n=118 mice, with retention interval of 24 hours to 1 month, **Fig. 1**). The first two PCs explaining 72% of the variance (57.7 and 14.3, respectively), were used to define the avoidance index (**Extended Data Fig. 2a-c**). These two PCs were conserved throughout the study and used for projecting other data. There were two axes of variation of data dispersion when projecting the mouse scores on the first two PCs. They approximately aligned onto the two diagonals (a weighted sum of PCA loadings yielded respectively: 0.35*PC1+0.27*PC2 and 0.31*PC1-0.33*PC2). For simplicity, we kept the diagonal axis {PC1–PC2} and {PC1+PC2}, and evaluated their respective weights by quantifying the steepness of their evolution between familiarization and retrieval (**Extended Data Fig. 2c**). The vector {PC1-PC2} gives weight to the number of detours, sniffings, side passages and U-turns while the vector {PC1+PC2} gives weight to the contact configuration and latency. The avoidance index was defined as follows:

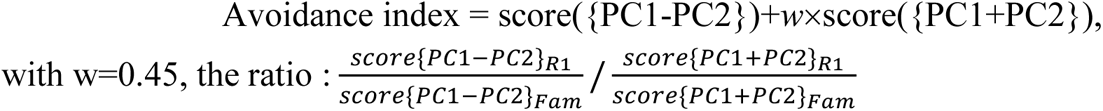

### Novel object recognition, rotarod and cross-maze tasks

See details and analysis in ***Supplementary Information***

### Viral injections

Viral injections were performed in S2 cortex of C57BL/6J mice, *CB1R^flox/flox^* and *Drd2^loxP/loxP^* mice (P35-40 days).

#### Viral vectors

To express ChR2 in L5 pyramidal neurons, AAV5-CAMKII-ChR2-mCherry (Addgene #26975, titer: 10^13^ particles/mL) was delivered unilaterally. To selectively ablate Drd2 or CB1 receptors in L5 pyramidal neurons, AAV1.CMV.HI.eGFP-Cre.WPRE.SV40 (Addgene #105545, titer: 10^13^ particles/mL, dilution 1/5) was delivered bilaterally in *Drd2^loxP/loxP^* or *CB1R^flox/flox^* mice. Injection of AAV1.CMV.HI.eGFP. WPRE.SV40 (Addgene #105530, titer: 10^13^ particles/mL, dilution 1/5) was used as a control.

#### Stereotaxic surgery

Two craniotomies were performed at the coordinates relative to bregma: {-3.85mm lateral, +0.5mm anterior, -3.05mm deep} and {-4.05mm lateral, -0.5mm posterior, -2.95mm deep}. AAV virus (350nL for each injection site) was injected with a 5µL Hamilton syringe, at a rate of 70-80nL/min. All *in vivo* behavioral and electrophysiological experiments or *in vitro* patch-clamp recordings were performed 3-5 weeks after viral injection.

### *In vivo* cortico-striatal ChR2-evoked LFP recordings and analyses

#### Optrode design and implantation

Two weeks following viral injection, a second surgery was performed to implant an optic fiber and custom electrical wires assembled to a 4-pin connector (Xiong et al., 2015). A stainless steel electrical wire (California Fine Wire, USA) composed of two strands, each soldered to one pin of the connector, was twisted and glued along a flat-cleaved 5mm length optic fiber of 0.39NA and 200μm core diameter (CFMLC 12L05, Thorlabs, Maisons Laffitte, France). The two strands were separated at the extremity of the optic fiber such that their tips reached the same height (±200μm) as the optic fiber. The optrode was implanted in DLS at (-2.9mm lateral; -0.1mm posterior; 3.7-3.9mm deep). Two miniature screws (Antrin Miniature Specialties Inc., Fallbrook, CA, USA) for ground and reference were screwed over the cerebellum. The implant was cemented to the skull using Superbond C&B dental cement (Sun Medical, Moriyama, Japan). Self-curing acrylic (Dentalon Plus, Kulzer) was used to cover the rest of the wires and the optic fiber, as well as to attach the skin. A copper mesh circlet was attached to the skull, protecting the implant.

#### Electrophysiological recordings

Extracellular signals were amplified, multiplexed and acquired continuously at 40kHz using a multi-channel KJE-1001 system (Amplipex, Szeged, Hungary). ChR2 was activated using a 475nm-laser diode light source (FLS-475, DIPSI, Cancale, France). The brain-implanted optic fiber cannula (ceramic ferrule of 1.25mm diameter and 6.4mm length) was connected to a FC/PC patch cable with similar ferrule diameter (ThorLabs). ChR2-LFP were recorded 10 minutes before, 2 hours and up to 4 hours after each training session in awake mice. Light was delivered for 10 minutes at 0.14Hz in square pulses of 300μs in average (range: 150-800μs) with a light power of 7-15mW at the tip of the fiber, controlled by an Arduino UNO.

#### Data analysis

Data were analyzed in MATLAB. For each light stimulation, the ChR2-LFP was extracted and conserved only if the slope of the first component was clearly identified. Each ChR2-LFP was smoothed (moving mean, window of 150μs) and realigned to zero by subtracting the average voltage of the 375μs preceding light stimulation. Each measurement of ChR2-LFP was obtained from averaging six successive conserved responses. The amplitude the second phase was normalized to the first phase amplitude. The significance of synaptic changes for each individual mouse was estimated using Repeated-Measures ANOVA and Tukey post-hoc tests, based on the 10-15 batch measurements obtained on each recording session.

#### In vivo and in vitro pharmacological characterization of cortico-striatal ChR2-LFP

*In vivo* pharmacology was performed via cannula-equipped mice (see below: canula implants), with infusion of 1μM TTX (Tocris, Ellisville, MO, USA) diluted in saline, combined with ChR2-LFP recordings. *In vitro*, the tip of the optic fiber was inserted at the surface of horizontal slices in DLS (see below: in vitro patch-clamp recordings). LFP, sampled at 20kHz, were recorded (EPC10, HEKA Elektronik, Lambrecht, Germany) using a glass pipette of 3-5MΩ filled with ACSF, positioned within the area illuminated by the optic fiber. Baseline recordings were performed under ACSF perfusion for 10 minutes. Then, various mixes of drugs diluted in ACSF perfused the slice successively, with 20min recording between each: D-AP5 (50μM), D-AP5+CNQX (20μM), D-AP5+CNQX+TTX (1μM) (Tocris).

### Cannulas implants and *in vivo* microinjections

Double guide cannulas (33 GA, Bilaney, Düsseldorf, Germany) were implanted per hemisphere at (+0.55mm anterior; -2.9mm lateral) and (-0.4mm posterior; -3.1mm lateral) at 2mm from the skull surface. Guide cannula dived down to 0.5mm from the skull surface. Double 33 G internal cannula (Bilaney, Düsseldorf, Germany) of 3.5mm length, with 2mm projection from the guide cannula, were inserted. After anesthesia removal, bilateral infusion was performed using 10μL Hamilton syringes link to automatic infusion pump. 40-50 minutes separated the infusion to behavioral test. 500μM D-AP5 diluted in saline were injected bilaterally (1μL per site at 0.1μL/min) into DLS.

### Neuropixel electrophysiological recordings

To perform acute Neuropixel recordings, we placed head-fixed mice in an air-lifted low-weight carbon cage (65g, 290mm diameter) using the Mobile HomeCage (MHC) (Neurotar, Helsinki, Finland).

#### Headplate implantation

Mice (P40-45 days) were head-fixed to the MHC after fixating a steel rectangular headplate (dimensions: 5,2mmx7,2mm, 1mm thick, weight: 1g, Neurotar), glued (VetBond, WPI) and cemented to the skull over the parietal and interparietal bone. A miniature screw (Antrin) was inserted on the anterior bone above the cerebellum and connected to a 2-pin connector, cemented on the headplate.

#### Neuropixel recordings

During brief anesthesia with isoflurane (1.5% maintenance, 100% O_2_), one craniotomy over S2 cortex was made in the left hemisphere (-0.1 mm posterior; -2.9 mm lateral). Dura was removed and the craniotomy was covered with Kwik Cast (WPI, USA). The mouse was head-fixed in the MHC and Neuropixel Phase 3b probe^61^, mounted on a 3-axis micromanipulator, was inserted at 10μm/s (Luigs & Neumann, Ratingen, Germany) to a final depth of 3.8-4mm. To allow track localization, the probe was coated with DiI. Signals, acquired using SpikeGLX (HHMI/Janelia Research Campus, Ashburn, VA, USA), were split into a spike band (30kHz sampling rate, 500Hz high-pass filter, gain: 500x) and an LFP band (2.5kHz sampling rate, 1000Hz low-pass filter, gain: 250x). The probe was connected to a PXIe card (PXI-e8381, National Instruments, Austin, TX, USA). The tip of the electrode was chosen as reference. Two to three weeks of mouse habituation to the MHC (see below) were allowed preceding the recording day. During 15-minute baseline recordings, the head-fixed mouse was left in the empty arena. Then, the mouse was subjected to the STA familiarization and recordings were performed up to 30 minutes after familiarization.

#### Data analysis

Spike sorting was performed offline using Kilosort 2.5^62,63^ and manually curated with the phy GUI interface (https://github.com/kwikteam/phy). Preprocessing performed in Kilosort 2.5 included high-pass filtering (300Hz), common average referencing to remove artefacts, and whitening. Each detected cluster containing at least 50 spikes was manually checked: if the template waveform was too close to noise (near-zero amplitude) or was non-physiologically realistic, the cluster was discarded. Units were then classified depending on their amplitude and signal-to-noise ratio, and the degree of refractory period contamination and merges were checked. Double-counted spikes due to templates fitting residuals from large-amplitude spikes were removed in a post-processing step, using a temporal and spatial window of 0.16ms and 50μm. Further analyses were performed on units with no/almost absent refractory period contamination and for which the spike amplitude distribution was not truncated by the detection threshold.

#### Mouse habituation to MHC

The mouse was habituated to the MHC for two to three weeks. On the first day, mice were left 5 min inside the cage, with the air pressure on, without head-fixation. Then, mice were head-fixated (with a 20° tilted head-clamp) and left for 15 minutes. Training duration progressively increased (on the first three days: 15, 30, 45 minutes, then on the subsequent days: alternation between 1 and 2 hours of training), with one session per day. Starting on the 5^th^ session, small objects were placed in the border or center of the cage forming obstacles to improve the anti-collision abilities of mice. After 8-10 sessions, headplate position relative to the MHC head-fixation bar was rotated by 90°. This position was preferred over time since mice better followed the natural circular movement of the cage and anticipated collisions with objects. Mice were also habituated to the presence of the electrophysiological apparatus and light (to mimic the light needed for probe insertion). On the day preceding the Neuropixel recordings, a special floor made of corrugated paperboard was added, which allowed a finer grip.

### *Ex vivo* and *in vitro* whole-cell patch-clamp recordings and analyses

#### Brain slice preparations

We used horizontal and para-sagittal (30°) brain slices to stimulate cortical inputs originating either from somatosensory or from the secondary anterior cingulate cortex, and projecting to DLS or DMS, respectively^24^. Brain slices (270µm-thick for DLS and 30µm-thick for DMS) were prepared with vibrating blade microtome (7000smz-2, Campden Instruments Ltd., UK) in a 95% O_2_ / 5% CO_2_-bubbled, ice-cold cutting artificial cerebrospinal fluid (ACSF) solution containing (in mM): 125 NaCl, 2.5 KCl, 25 glucose, 25 NaHCO_3_, 1.25 NaH_2_PO_4_, 2 CaCl_2_, 1 MgCl_2_.

#### SPN whole-cell patch-clamp recordings

Borosilicate glass pipettes of 4-7MΩ resistance were filled with (in mM): 122 K-gluconate, 13 KCl, 10 HEPES, 10 phosphocreatine, 4 Mg-ATP, 0.3 Na-GTP, 0.3 EGTA (adjusted to pH 7.35 with KOH). The extracellular solution was the same ACSF solution used for brain slice incubation through which 95% O_2_ and 5% CO_2_ was bubbled. Signals were amplified using EPC9-2 amplifiers (HEKA Elektronik, Lambrecht, Germany). All recordings were performed at 34°C (Bath-controller V, Luigs&Neumann, Ratingen, Germany). Recordings were sampled at 10kHz, using the Patchmaster v2x32 program (HEKA Elektronik).

#### Spike-timing dependent plasticity (STDP): protocols and off-line analysis

Electrical stimulations were performed with concentric bipolar electrodes (Phymep, Paris, France) placed in cortical layer 5. Electrical stimulations were monophasic, at constant current (ISO-Flex stimulator, AMPI, Jerusalem, Israel). Currents were adjusted to evoke 50-300pA EPSCs at 0.1Hz. STDP protocols consisted of 15 pairings of post-presynaptic stimulations (at 1 or 2.5Hz) with Δt_STDP_=-15ms)^30^. Postsynaptic stimulation corresponded to two action potentials evoked by a depolarizing current step (30ms duration) in the recorded SPN (hold at -70mV), and presynaptic stimulations to cortical stimulations. Recordings were made over a period of 10 minutes at baseline, and for at least 30 minutes after the STDP protocol. Input and series resistances were monitored during each sweep and experiments were excluded if these parameters varied by more than 20%. Off-line analysis was performed with custom MATLAB code. In all cases “n” refers to a single-cell experiment from a single brain slice.

#### Occlusion/saturation ex vivo experiments

Mice subjected to no-tape and STA tests were divided into two groups for occlusion/saturation experiments: (1) post-familiarization and (2) post-retrieval mice. After task completion, mice were left in their homecage for 30 minutes and then sacrificed for *ex vivo* whole-cell recordings. Plasticity occlusion/saturation was tested with 15 post-pre pairings (1Hz, Δt_STDP_ =-15ms).

### Histology

Injection sites and extent of virus expression were verified for every animal. Brains were post-fixed in PFA (4% in PBS) and then cryoprotected in 30% sucrose solution. Brains were cut in coronal 80µm sections, maintained in 0.1M potassium-PBS (pH=7.4). Slices were stained with DAPI and mounted on glass slides, coverslipped with Fluoromount (Thermofisher, Asnières-sur-Seine, France). Images were acquired using a stereozoom fluorescence microscope (Axiozoom, Zeiss, Germany) and processed in ImageJ.

### Theoretical tools for plasticity induction estimation

#### Mathematical model

The potential for expression of synaptic plasticity was estimated using a mathematical model that describes the effect of input spike trains on the concentration of the main signaling actors involved in STDP at the cortico-striatal synapse in mice^31^. The model expresses the kinetics of the involved molecular reaction pathways and ionic channel behaviors at a biophysical level through non-linear ordinary differential equations. The signaling partners included in the model comprise e.g., glutamate receptors, VSCC, CB1R or IP3R channel receptors, enzymes including FAAH, CaMKII kinase or DAG lipase, and smaller metabolites, molecules or ions including glutamate, Ca^,+^, DAG, 2-AG or IP_3_. The model uses the ratio of activated (phosphorylated) CaMKII in the postsynaptic terminal as a proxy for NMDAR-dependent spike-timing dependent LTP. The level of CB1R activation by endocannabinoids (2-AG and anandamide) in the presynaptic bouton is used as a proxy for endocannabinoid-dependent plasticity, with the following experimentally-validated rule: intermediate levels of CB1R activation triggers eCB-LTD whereas large levels of CB1R activation give rise to eCB-LTP^31^. This model has been extensively calibrated and validated ex-vivo on mouse slice STDP experiments^12,29,31^. Extensions of this model were developed and validated in a range of different experimental contexts^30,37^. Here, we used the original version of the model without modification^31^ in the equations or parameter value, except for the glutamate release per pre-synaptic spike parameter *Glu*_max_ and the duration of the postsynaptic depolarization associated to each postsynaptic spike *DP*_dur_, that we altered to account for the difference in activity levels between *in vivo* and *in vitro* situations. In slice experiments, presynaptic stimulations induces a large amount of synchronized presynaptic input while spike sorting of *in vivo* recording gives access to single-cell spike trains. To account for this difference, we decreased *Glu*_max_ to 1.0 mM. In the slice STDP experiments used to calibrate the model, post-synaptic stimulation consisted of a 30ms depolarization on which the spike seats. While population effects in the striatum are likely to give rise to similar depolarization events around each postsynaptic spike, we decreased their duration to 14ms.

#### Numerical simulation

The cerebral area of each sorted neuron was deduced from the position of the corresponding individual electrode along the Neuropixel probe, allowing to classify the individual activities as cortical or striatal neurons. Each pair of cortical and striatal neurons was evaluated for potential cortico-striatal plasticity expression, using the experimental spike trains as inputs to the model. Simulations were performed using a custom code in Python 3.10.12 using Runge-Kutta 4-5 (“RK45”) solver from the scipy package^64^, and optimized with numba^65^. Data management was supported by the Hydronaut package (https://gitlab.inria.fr/jrye/hydronaut). The model was used to evaluate plasticity expression starting from 5 minutes before contact time up to 5 minutes after contact time. Code is freely available at: https://gitlab.inria.fr/aistrosight/ecb_ltp_oneshotlearning_paper.

#### The plasticitymeter

The original mathematical model was calibrated on short (around 1 minute) and non-bursty stimulation typical of STDP protocols in brain slices. In contrast, *in vivo* spike trains recorded here are much longer (>10 min) and more bursty. We therefore needed a robust indicator over the course of the recording but fine enough to resolve local induction of plasticity around the contact. In the original mathematical model, the presynaptic weight *w*_pre_ is estimated from the amount of calcium and the level of CB1R bound to endocannabinoids like 2-AG, according to *dw*_pre_ ∕ *dt* = (Ω − *w_pre_*) ∕ σ, where σ is a time scale that decreases with increasing calcium concentration and defines the delay for full expression of the plasticity in *ex vivo* slice experiments, Ω is a CB1R-dependent function that defines the *w*_pre_ variation expected from the current synaptic inputs. Ω < 0 when the level of 2-AG binding to CB1R is intermediate, thus leading to LTD, whereas Ω > 0 when the level of 2-AG binding is large, leading to LTP^31^. We combined the values of Ω and σ into a robust quantifier of the local triggering of plasticity expression that we refer to as the “plasticitymeter”. We first binned the spike trains into *N* consecutive non-overlapping temporal bins of 0.2 second. In each 0.2 second bin, we set the plasticitymeter of the pair (*i,j*), *π_i,j_* to 1 if at least one plasticity event occurred within the bin, and 0 else. We defined a plasticity event as a simultaneous occurrence of Ω in the LTP state (eCB-LTP event), or in the LTD state (eCB-LTD event), together with values of σ low enough to allow some maintenance of the weight change (we used an arbitrary threshold of σ < 100). We kept separate counts of eCB-LTP and eCB-LTD events.

We then defined the plasticitymeter of a sub-population *A* of pairs, as the mean of normalized plasticitymeter over pairs. The plasticitymeter *P* of sub-population *A* at bin *k* is given by:

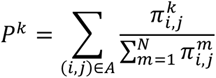

We used the temporal mean μ = 1/*N* ∑*_k_ P*^k^ and corresponding standard deviation *s* of the plasticitymeter values to obtain its z-score *z_k_* = (*P^k^* − μ) ∕ *s*.

#### Sub-selection of neuron pairs

Even with the above plasticitymeter algorithm, the quantity of possible cortical-striatal pairs makes it difficult to analyse the Neuropixel data. We therefore searched to focus the analyses on pairs that are more likely to be connected together. We quantified the degree of correlation of each neuron pairs (*i*, *j*) during the contact with the sticky tape:

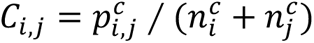

where:

- 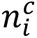 is the number of spikes in the train of neuron *i*, during the first 20 seconds of contact or the last 290 seconds of baseline, 10 seconds before the contact, respectively
- 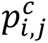 is the number of pre-post or post-pre spike pairs separated by 0.2 seconds or less, during the contact or baseline epochs defined as above.

The neuron pairs (*i*, *j*) with a *C_i,j_* within the top 5% formed the *active neuron (contact)* group, **Fig. 3e**, where we show, for each 0.2 second time bin, the z-score of the neuron-pairs of the active neuron (contact) group.

### Statistics

Results are expressed as mean ± SEM. Statistical significance was assessed using MATLAB or Jamovi (http://www.jamovi.org). All two-sample and one-sample tests were two-tailed. For **Fig. 1: b**, Linear Mixed Model for STA (n=36, 34, 19) and n-STA (n=22) between zones (close to the tape location in yellow, 90° and 180° opposite wall locations in green and red respectively) and between the two groups. Cluster variable corresponded to mouse ID. Levels of statistical significance are similar for all three timepoints. For occupation probability. Fixed Effects of {Group: *F*=8.822; *p*=0.003; Timepoints *F*=8.20, *p*<0.001; Zone: *F*=6.86, *p*=0.001 and Group*Zone: *F*=17.1, *p*<0.001}. For average speed. Fixed Effects of {Group: F=1.26; *p*=0.263; Timepoints: *F*=27.7, *p*<0.001; Zone: *F*=30.3, *p*=0.001 and Cluster*Zone: *F*=14.8, p<0.001}. Post-hoc Tukey tests Cluster*Zone with Bonferroni corrections: STA Scotch/90°: p<0.001; nSTA Scotch/90°: p=1.00 for both occupation probability and speed. (**c**) Comparison of behavioral features between STA (grey, n=56 mice) and n-STA (blue, n=22 mice) for familiarization and retrieval. (**c**) Comparison of behavioral features between STA (grey, n=56 mice) and n-STA (blue, n=22 mice) for familiarization and retrieval. **c**, Contingency Tables: between retrieval ≥1 detour: χ^2^=21.1, *p<*0.001 and ≥2 detours: χ^2^=18.1, *p<*0.001; between familiarization ≥1 detour: χ^2^=0.414, *p=*0.520 and ≥2 detours: χ^2^=0.084 *p=*0.771. McNemar-Test for paired samples: within n-STA for ≥1 detour: χ^2^=0.143; *p=*0.705 and ≥2 detours: χ^2^=18, *p<*0.001; within STA for ≥1 detour: χ^2^=23.1, *p<*0.001 and ≥2 detours: χ^2^=11.6, *p<*0.001. **d**, Repeated-Measures ANOVA. Between-Subjects: *F=*16.2, *p<*0.001; familiarization-retrieval: *F=*17, *p<*0.001; Combined: *F=*16.3, *p<*0.001. Post-hoc Tukey tests with Bonferroni correction: STA familiarization/retrieval: *t=*-7.7, *p<*0.001; nSTA familiarization/retrieval: *t=*-0.0482, *p=*1.0; familiarization nSTA/STA: *t=*0.0209, *p=*1.0; retrieval nSTA/STA: *t=*4.069, *p<*0.001. **e**, Repeated-Measures ANOVA. Between-Subjects: *F=*13.1, *p<*0.001; familiarization-retrieval: *F=*16, *p<*0.001; Combined: *F=*13.4 *p<*0.001. Post-hoc Tukey tests with Bonferroni correction: STA familiarization/retrieval: *t=*-7.2, *p<*0.001; nSTA familiarization/retrieval: *t=*-0.197, *p=*0.997; familiarization nSTA/STA: *t=*0.281, *p=*0.992; retrieval nSTA/STA: *t=*3.673, *p=*0.002. **f**, Log-rank test (Mantel-Haenszel Hazard ratio, MHZ): within n-STA: MHZ=1.14; *p=*0.79; within STA: MHZ=5.74; *p<*0.001; between familiarization: MHZ=1.28; *p=*0.47; between retrieval: MHZ=8.36; *p<*0.001. **g**, Paired two-sample t-test: *t=*3.23; *p=*0.002. **h**, Paired two-sample t-test: *t=*-10.4; *p<*0.001. McNemar test for avoider rate: χ^2^=30.1, *p<*0.001. k, χ^2^-test on avoider rate: χ^2^=6.14, *p=*0.105 for contact duration (right); χ^2^=4.71, *p=*0.194 for time-to-remove (left). **l**, One-way ANOVA. Latency-to-contact: *F=*2.093, *p=*0.101; number of detours: *F=*0.430, *p=*0.786. Two-sample t-test with or without extra-task learning (latency-to-contact: *t=*0.364, *p=*0.718; detours: *t=*0.142, *p=*0.888). **m**, One-Way ANOVA on avoidance index: *F=*1.051; *p=*0.394. Two-sample t-test between 1 month with/without extra-task: *t=*0.366, *p=*0.717. χ^2^-test on avoider rate across all conditions: χ^2^=0.877, *p=*0.928.

For **Fig. 2**: **b**, Linear Mixed Model: STA (n=23): *F=*4.74, *p=*0.002; no tape (n=11): *F=*3.73, *p=*0.011; n-STA (n=7): *F=*0.559, *p=*0.695. **c**, One-Way ANOVA: *F=*1.02, *p=*0.415. One-sample t-test for LTP significance: no tape: *t=*0.351, *p=*0.733; n-STA: *t=*0.561, *p=*0.596; STA: *t=*2.44, *p=*0.0229. **d**, Two-sample t-test: *t=*-2.33; *p=*0.0295. **e**, Linear Mixed Model with effect of contact duration (*F=*12.10; *p=*0.002), time (*F=*3.97; *p=*0.006) and combined (*F=*4.55, *p=*0.003). **f**, One-sampled t-test for LTP significance on each data point. ANCOVA on avoidance with two control binary variables {significant LTP and contact duration (≤20 seconds)}: no effect of each control variable (*F=*0.006 and *F=*0.586 respectively), effect of combined control variables (*F=*4.83, *p=*0.041).

For **Fig. 3**: **b**, Reference for baseline firing rate is the 90^th^ percentile of the average firing rates found in the same contact time window among 310s to 10s preceding contact with tape. Ratio of mean firing rate during contact over baseline reference. Two-sample t-test for cortex and striatum, respectively: ID#1 (*t*=-11.45, *p*<0.001; *t* = -0.48, *p*=0.631), ID#2 (*t* =-9.63, p<0.001; *t*=1.56, *p*= 0.123), ID#3 (t=0.33, *p*=0.741; *t*=3.52, *p*=0.001), ID#4 (*t*=0.62, *p=*0.539; *t*=-2.68, *p*=0.013). **e**, z-scores of the plasticitymeter for active pairs are > 2 for all mice in at least 10% of the time bins within the first 10 seconds after contact onset; in n=3 mice (exluding ID#2), scores of the plasticitymeter for active pairs are > 6 in at least 10% of the time bins within the first 20 seconds after contact onset (> 3.5 in 20% time bins). For all mice, the proportion of z-scores>2 during the 20 seconds from contact onset is at least 3 times larger than during the 5min baseline preceding contact; excluding ID#2, it becomes at least 4 times larger. **f**, One sample Wilcoxon test (against 100%). Post-familiarization: W=91, *p<*0.001 for no-tape (n=13); W=20, *p=*0.375 for STA (n=7). Post retrieval: W=66, *p<*0.001 for no tape (n=11); W=0.5614, *p=*0.561 for STA (n=15). Two-sample Wilcoxon test. Post-familiarization: W=170, *p=*0.0089; post retrieval: W=214, *p*<0.001.

For **Fig. 4**: **b**,**g**, Two-sample Wilcoxon test between (**b**) CB1R-Cre (n=10) and CB1R-ctrl (n=9): W=118, *p=*0.022; (**h**) Drd2-Cre (n=8) and Drd2-ctrl (n=9): W=114; *p*=0.0012. One sample Wilcoxon test (against 100%) for CB1R-Cre: W=39; *p=*0.2754; CB1R-ctrl: W=45; *p=*0.0039; Drd2-Cre: W=5, *p=*0.1562; Drd2-ctrl: W=44, *p=*0.0078. **c,i**, Two-sample Welch-test (for short contact duration, ≤30 seconds, because of unequal variance) and Student t-test (long contact duration >30 seconds) for latency-to-contact and detours, respectively. CB1R^flox/flox^ (**c**): CB1 short (n=6 Cre and n=10 ctrl): *t=*3.58, *p=*0.006; *t=*2.75, *p=*0.022; CB1 long (n=5 Cre and n=9 ctrl): *t=*1.25, *p=*0.235; *t=*1.48, *p=*0.166. Drd2^LoxP/LoxP^ (**i**): D2 short (n=9 Cre and n=13 ctrl): *t=*2.76, *p=*0.015; *t=*2.47, *p=*0.027; D2 long (n=10 Cre and n=8 ctrl): *t=*-0.784, *p=*0.445; *t=*-0.718, *p=*0.483. **d**,**i,** Two-sample Welch (for short contact because of unequal variance) and Student t-test (long) for avoidance index. CB1R^flox/flox^ (**d**): CB1 short: *t=*3.26, *p=*0.008; CB1 long: *t=*1.18, *p=*0.262. Drd2^LoxP/LoxP^ (**i**): D2 short: *t=*3.09; *p=*0.006; D2 long: *t=*-1.39; *p=*0.182. χ^2^-test on avoider rate. CB1R^flox/flox^ (**d**): CB1 short: χ^2^=5.76, *p=*0.016; CB1 long: χ^2^=1.66, *p=*0.198. Drd2^LoxP/LoxP^ (**i**): D2 short: χ^2^=5.59, *p=*0.018; D2 long: χ^2^=3.38, *p=*0.066 (**i**). **e**,**j,** Linear Mixed Model for short contact only. CB1R^flox/flox^ (**e**): Fixed Effect of {Group: *F=*5.24, *p=*0.038; Time: *F=*5.69, *p=*0.008; Group*Time: *F=*3.29, *p*=0.052}. Fixed Effect of Parameter Estimates {Group: *t=*-2.289, *p=*0.038; retrieval-1-familiarization: *t=*2.307, *p=*0.029; retrieval-2-familiarization: *t=*3.285, *p=*0.003; Group*(retrieval-1-familiarization): *t=*-2.289, *p=*0.018; Group*(retrieval-2-familiarization): *t=*-0.859; *p=*0.398}. Drd2^LoxP/LoxP^ (**j**): Fixed Effect of {Group: *F=*13.44, *p<*0.001; Time: *F=*13.30, *p<*0.001; Group*Time: *F=*2.53, *p*=0.089}. Fixed Effect of Parameter Estimates {Group: *t=*-3.666, *p<*0.001; retrieval-1-familiarization: *t=*3.0247, *p=*0.004; retrieval-2-familiarization: *t=*5.124, *p<*0.001; Group*(retrieval-1-familiarization): *t=*-2.152, *p=*0.036; Group*(retrieval-1-familiarization): *t=*-0.490; *p=*0.626}. **l**, Two-sample t-test for latency-to-contact and detours. For short contact (n=7 D-AP5 and n=5 saline): *t=*0.297, *p=*0.773 and *t=*1.283, *p=*0.229 respectively; for long contact (n=12 D-AP5 and n=3 saline): *t=*-0.866, *p=*0.402 and *t=*-0.397, *p=*0.698. **m**, Two-sample t-test for avoidance index: *t=*1.11, *p=*0.292; *t=*-1.29, *p=*0.219 for short and long contact, respectively. χ^2^-test on avoider rate: χ^2^=1.66, *p=*0.198; χ^2^=0.417, *p=*0.519 for short and long contact, respectively. **n,** Linear Mixed Model for short contact only. Fixed Effect of {Group: *F=*0.657, *p=*0.437; Time: *F=*20.387, *p<*0.001; Group*Time: *F=*0.466, *p*=0.634}. Fixed Effect of Parameter Estimates {Group: *t=*-0.810, *p=*0.437; retrieval-1-familiarization: *t=*4.667, *p<*0.001; retrieval-2-familiarization: *t=*6.108, *p<*0.001; Group*(retrieval-1-familiarization): *t=*-0.662; *p=*0.516; Group*(retrieval-2-familiarization): *t=*0.278; *p=*0.784}.

## Acknowledgements

We thank S. R. Datta and the Venance lab members for helpful suggestions and critical comments on the manuscript. Camille Chataing and Emma Idzikowkski for their help on the behavioral experiments at one-month retrieval interval. Yves Dupraz (CIRB micromechanics workshop) for the building of the arenas, cross-maze and electrophysiology micromechanics.

## Author contributions

Conceptualization: L.V., C.P. and J.T. C.P. performed behavioral experiments, viral injections and *in vivo* recordings; C.P. performed analysis of the behavioral and electrophysiology experiments; C.P. and S.P. performed patch-clamp experiments and surgeries for *in vivo* opto-LFP experiments and canula implants; C.P. and A.H. performed spike sorting of Neuropixel recordings; A.H. and H.B carried out the conception and the design of the mathematical model; A. H. and H.B. performed the acquisition and analysis of data from the mathematical model; C.P and L.V. wrote the manuscript and all authors have edited and corrected the manuscript; Funding acquisition: L.V.; L.V. supervised the whole study.

## Competing interests

The authors declare no competing interests.

**Supplementary Information is available for this paper.**

## Extended Data Figures (from 1 to 10)

**Extended Data Figure 1:**
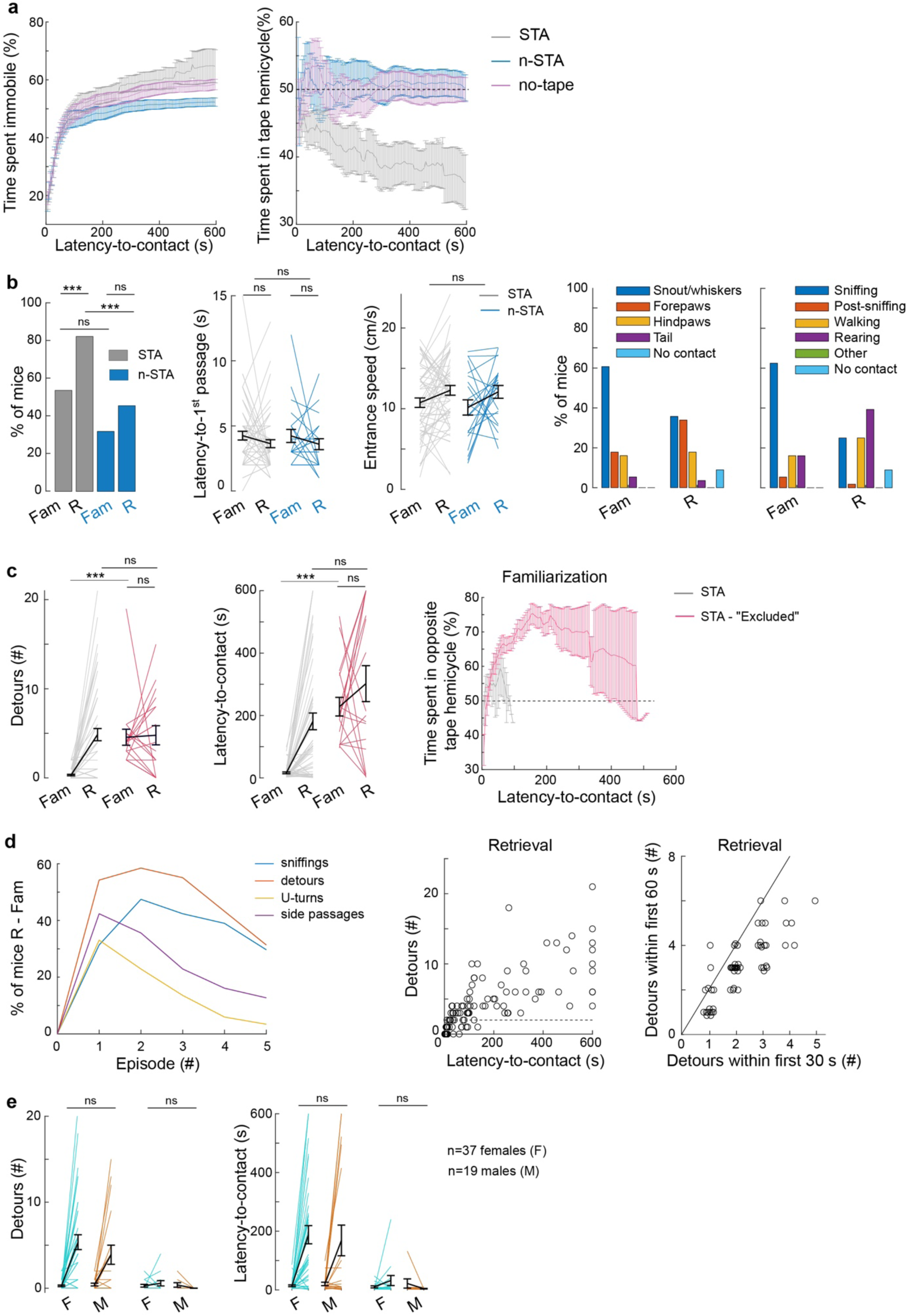
Characterization of the STA and n-STA tests. **a**, Time spent immobile (right) or in the hemicycle arena containing the tape (or randomly centered in no-tape condition) (left) up to the latency-to-contact show an avoidance behavior at retrieval in STA mice but not in mice exposed to n-STA and no-tape conditions. Linear Mixed Model. For time spent immobile: fixed effect of groups: *F=*1.022, *p=*0.036; timepoints: *F=*0.125, *p<*0.001 and groups*timepoints: *F=*0.773, *p*=0.996; for time spent in the hemicycle arena containing the tape: fixed effects of groups: *F=*8.02, *p<*0.001; timepoints: *F=*4.09, *p<*0.001 and groups*timepoints: *F=*0.675, *p=*1.000. **b**, Left: The non-repelling nature of the tape is indicated by the percentage of mice having at least one passage near the tape before contact. Between familiarization: χ^2^=3.0, *p=*0.086; Between retrieval: χ^2^=10.5, *p=*0.001. McNemar-Test for paired samples: within n-STA for ≥1 passage: χ^2^=1.92 *p=*0.166; within STA: χ^2^=13.3 *p<*0.001. Middle: latency-to-1^st^ passage near the tape and entrance speed do not differ between n-STA and STA mice. Repeated-Measures ANOVA: between subjects: latency first passage *F=*0.381; *p=*0.539 (latency-to-first passage); *F=*0.198, *p=*0.657 (entrance speed); within subjects (familiarization *vs*. retrieval): *F=*0.747, *p=*0.390 (latency-to-first passage); *F=*5.87, *p=*0.018 (entrance speed). Right: Distribution of body part initially contacting the tape and activity preceding contact between familiarization and retrieval in STA mice. Log-linear regression model. For body part sticking to the tape (R^2^=0.843), familiarization/retrieval predictor: *p=*1.000; for activity immediately pre-contact (R^2^=1), familiarization/retrieval predictor: *p=*0.004. **c**, In STA, a subset of mice (24%, n=12 females, n=6 males) shows neophobic-like behavior at familiarization (red). Left: number of detours and latency-to-contact were larger than the rest of the mice (grey) at familiarization while they did not differ significantly at retrieval. Repeated-Measures ANOVA: between subjects: F=8.71, p=0.004 (detours) and F=30.9; *p*<0.001 (latency-to-contact). Right: Evolution of time spent in the hemicycle arena opposite to the sticky tape location. Comparison to a 50-50% occupancy at time +30 and +60 seconds: *t=*-3.1, *p=*0.0063; *t=*-5.9, *p<*0.001 for avoiding mice; *t=*-0.7, *p=*0.51 and *t=*-1.2, *p=*0.32 for the rest of mice. **d**, Left: Difference in the proportion of mice exhibiting a given number of sniffings, detours, U-turns or side passages between retrieval and familiarization. The number for which the maximal difference was obtained later served as a truncation threshold for defining the avoidance index. Right: Evolution of the number of detours as a function of the latency-to-contact at retrieval, and zoom on the sublinear relationship between the number of detours within the first 30 and 60 seconds after entering the arena at retrieval. **e**, Males (STA: n=19, n-STA: n=8) and females (STA: n=36, n-STA: n=16) did not show difference in number of detours and latency-to-contact. One-Way ANOVA (only sex effect is indicated here for detours and latency-to-contact respectively): for n-STA: *F=*0.925, *p=*0.348; *F=*0.428; *p=*0.520; for STA: *F=*0.796, *p=*0.376; *F=*0.0537, *p=*0.818. All data: mean±SEM. \**p<*0.05; \*\**p<*0.01; \*\*\**p<*0.001.

**Extended Data Figure 2:**
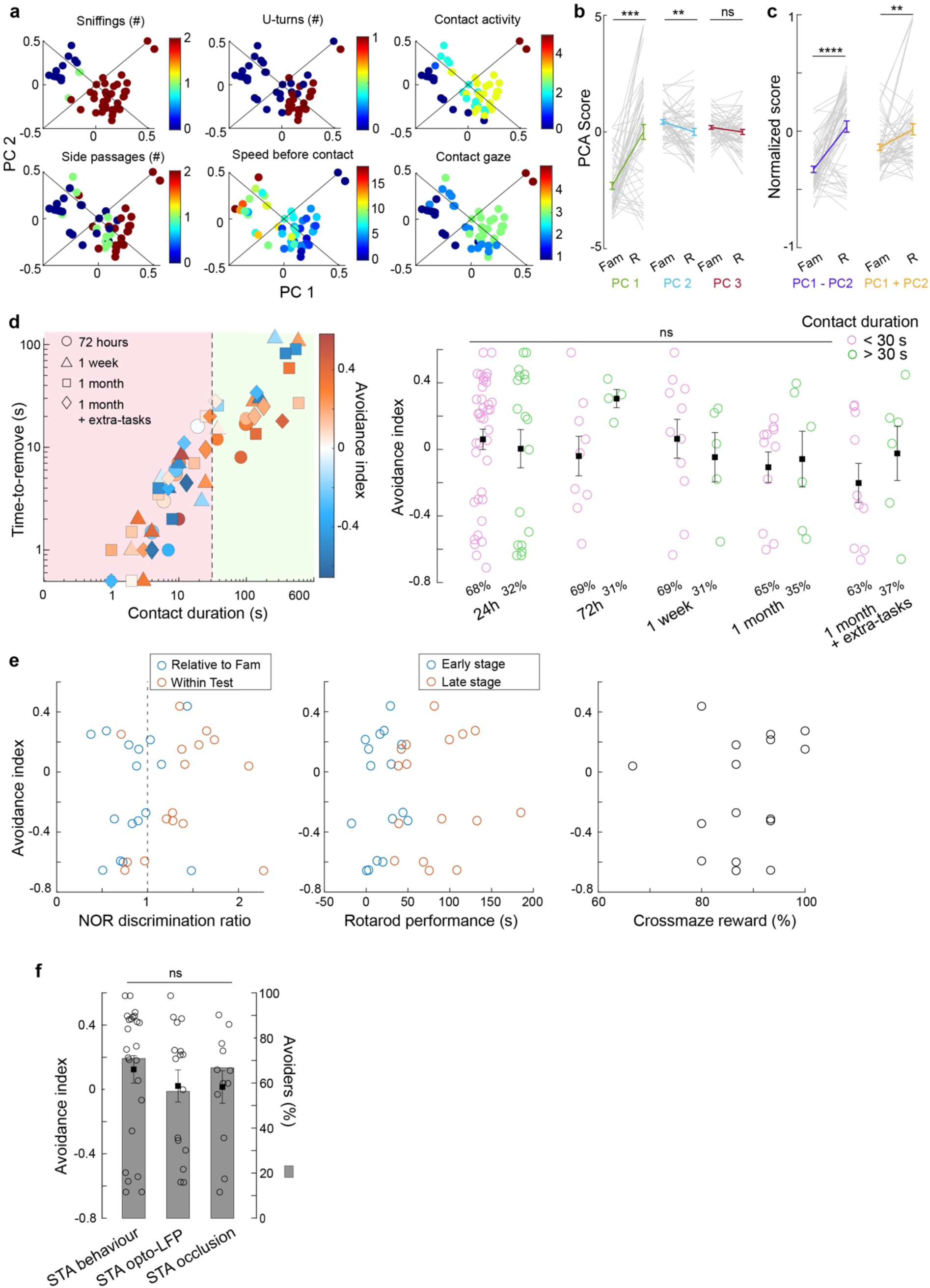
Evaluation of one-shot learning performance and its robustness. **a-c**, One-shot learning avoidance index. **a**, Six additional panels of retrieval behavioral parameter distribution in PCA space, similar to Figure 1i. **b**, Comparison of the individual mouse scores on the first three PCs between familiarization and retrieval shows significant differences for PC1 (green, *t=*-6.45, *p<*0.001) and PC2 (blue, *t=*3.31, *p=*0.002). For PC3 (*t=*0.088, *p=*0.93) (paired two-sample t-test). **c**, Two linear combinations of PC1 and PC2 {PC1-PC2} (purple) and {PC1+PC2} (yellow) capture most variance of the first two PC scores and show significant increases between familiarization and retrieval with different steepness (PC1-PC2: *t=*-7.0, *p<*0.001; PC1+PC2: *t=*-3.01, *p=*0.004) (paired two-sample t-test). The ratio of these two slopes, equal to 0.45, is used to weight the contribution of each linear combination in the avoidance index (*Methods*). **d**, Contact duration does not affect learning performance when retention intervals are introduced between familiarization and retrieval. Left: Avoidance index (color-coded) distribution as a function of contact duration and time-to-remove (log scale). The length of the retention interval is indicated with symbols: circle: 72 hours; triangle: 1 week; square: 1 month; diamonds: 1 month + extra-tasks. Right: Average avoidance index is not affected by contact duration (2-way ANOVA, contact duration: *F*=0.873, *p=*0.352; condition: *F=*0.908, *p=*0.462; intersection: *F=*0.863, *p=*0.489). Percentage of mice in each condition (contact duration ≤30 or >30 seconds) is indicated at the bottom. (**e**) In the configuration {1 month + extra-tasks}, STA performance does not correlate with learning performance of NOR, crossmaze and rotarod tasks. Left: NOR discrimination ratio: blue circles for time spent on the novel object at Test / average time spent on novel objects at familiarization; orange for time spent on the novel object at Test / average time spent on familiar objects at Test. Middle: Accelerating rotarod performance: the indices subtract average performance of the last 2 trials of Day 1 (early stage), or the last 2 trials of Day 5 (late stage) to the average of the first 2 trials of Day 1. *Right*: Crossmaze performance (% of rewarded trials on the last day of training). Correlations between avoidance index and NOR discrimination ratio relative to familiarization and within respectively (Pearson’s r=0.08 and 0.22; *p=*0.768 and 0.424 respectively); early and late rotarod performance (Pearson’s r=0.17 and 0.018; *p=*0.537 and 0.949 respectively); crossmaze performance (Pearson’s r=0.11, *p=*0.679). **f**, No difference in the avoidance index and avoiders (grey bars) at retrieval (24 hours from familiarization) were observed between mice used for STA behavior only (n=24), *in vivo* opto-LFP experiments (n=16) and *ex vivo* occlusion (n=12). One-way ANOVA for avoidance index: *F*=0.46, *p=*0.6354; Chi-squared test for avoider rate: χ^2^=0.488 *p=*0.784. All data: mean±SEM. \**p<*0.05; \*\**p<*0.01; \*\*\**p<*0.001.

**Extended Data Figure 3:**
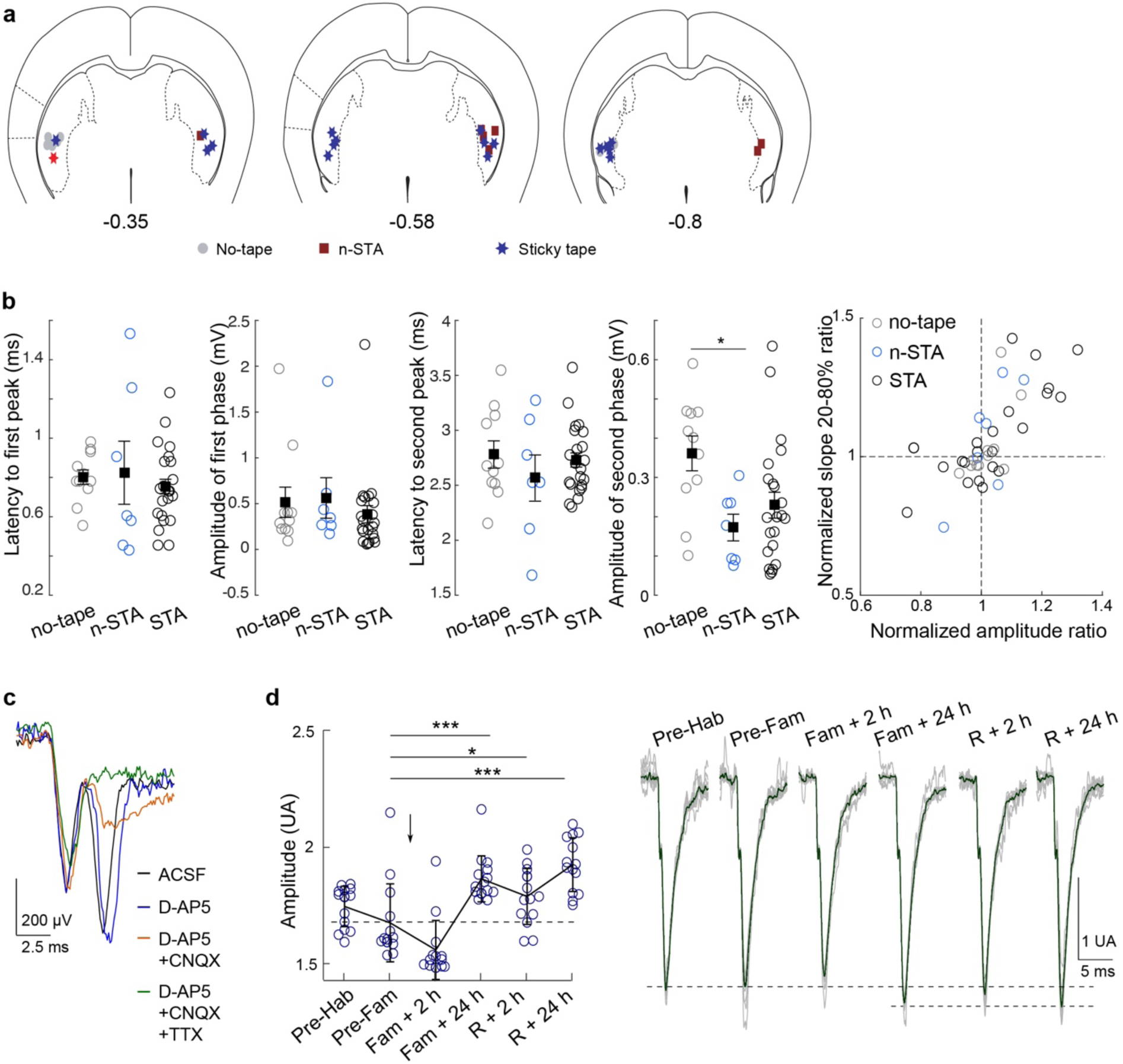
ChR2-LFP characterization. **a**, Recording sites in DLS are indicated on three coronal slices (0.3, 0.58 and 0.8 mm posterior to bregma). No-tape, n-STA and STA mice are represented with grey circles, red squares and blue stars, respectively. **b**, *Left*: Latency and amplitudes of the early and late ChR2-LFP phases across groups (One-way ANOVA: latency 1^st^: *F=*0.309, *p=*0.739; amplitude 1^st^: F=0.309, *p=*0.739; latency 2^nd^: *F=*0.368; *p=*0.699; amplitude 2^nd^: *F*=5.65, *p=*0.012). *Right*: Plasticity ratios obtained at familiarization + 24 hours, using the amplitude or slope (20-80%) of the average normalized response, are consistent (Pearson’s r=0.909, *p<*0.001). **c**, Pharmacological blockade *in vitro* shows a strong reduction of the late phase of ChR2-LFP under D-AP5 (50 mM) and CNQX (1 mM) and a complete abolishment under TTX (1 µM). **d**, Example of chronic recordings during STA. Left: amplitude (mean±SD) of the second phase (measured on the response normalized by the amplitude of the first phase). Each dot represents the amplitude on the average of 6 consecutive stimulations. One-way ANOVA: *F=*33.9, *p<*0.001 (post hoc Tukey tests). Right: Illustrative traces from the same mouse (grey traces indicate averages of 6 consecutive stimulations; dark trace is the total average). All data (except D): mean±SEM. \**p<*0.05; \*\**p<*0.01; \*\*\**p<*0.001.

**Extended Data Figure 4:**
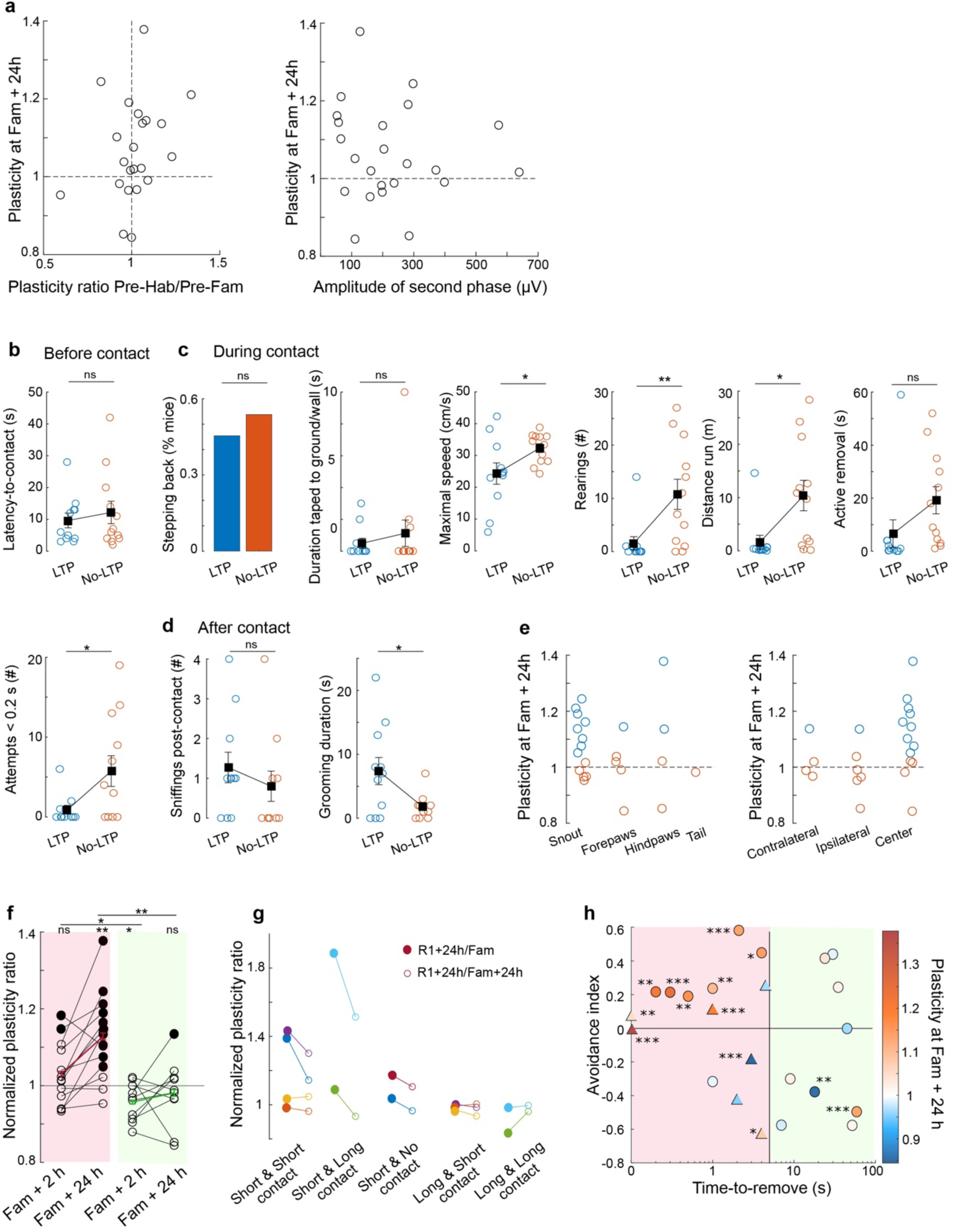
Relation between ChR2-LFP plasticity and behavioral features. **a**, Left: Plasticity ratio measured at familiarization + 24 hours as a function of the plasticity ratio between Pre-habituation/Pre-familiarization (informing about the expression of possible potentiation following first exposure to the arena at habituation, corresponding to a ratio < 1) does not present any significant correlation (Pearson’s r=0.286, *p=*0.1973). Right: Absence of correlation between plasticity expressed at familiarization + 24 hours and amplitude of second phase of ChR2-LFP response at familiarization (Pearson’s r=-0.1030, *p=*0.6399). **b-e**, Dependence of ChR2-LFP expression on behavioral parameters at familiarization (n=23 mice): (**b**) before contact, (**c**) during contact, and (**d**) after contact. Significant differences (two-sample independent t-tests) were observed during contact for the distance travelled (*p=*0.013), the maximal speed (*p=*0.032), the number of rearings (*p=*0.009), the number of very short (<0.2 second) attempts at removing the sticky tape (*p=*0.029) and after contact with the time spent in grooming 1 minute after contact (*p=*0.027). **e**, Left: Body part stuck to the sticky tape does not influence the induction of the ChR2-LFP potentiation (Chi-squared test, χ^2^=3.46, *p=*0.327). Right: Significant LTP was observed for all three contact positions relative to recording side (Chi-squared test, χ^2^=5.56, *p=*0.062). **f**, Plasticity ratio at familiarization +2 hours and familiarization + 24 hours for short and long contact duration (threshold ≤20 seconds). Significant LTPs are represented with filled circles. One-sample t-test: mice with short contact duration at familiarization +2 hours: {*t=*1.36, *p=*0.20}; short-familiarization +24 hours: {*t=*4.01, *p=*0.0017}; long contact duration at familiarization +2h: {*t=*-2.73, *p=*0.023}; long contact duration at familiarization +24 hours: {*t=*-0.70, *p=*0.50}. Two-sample t-tests between short and long contact duration at familiarization +2 hours: {*t=*2.52, *p=*0.020} and at familiarization + 24 hours: {*t=*3.38, *p=*0.0028}. **g**, Plasticity ratio measured at retrieval +24 hours normalized either by familiarization (filled) or familiarization +24 hours (empty) depending on the contact duration at familiarization and retrieval. The majority of mice showed stable plasticity profiles between familiarization +24 hours and retrieval +24 hours, and in particular mice (n=3) with long and then short contact duration at familiarization and retrieval respectively still did not express ChR2-LFP potentiation at retrieval +24 hours. **h**, Plasticity ratio at familiarization +24 hours (color-coded) as a function of STA avoidance index and time-to-remove. Circles and triangles represent mice with delays between familiarization and retrieval of 24 hours and longer (48 hours or 1 week), respectively. Statistical significance of each individual LTP is indicated (one-sampled t-test). ANCOVA on avoidance with two binary control variables {significant LTP and time-to-remove (≤5 seconds)}: no effect of each control variable (*F=*0.043 and *F=*1.61 respectively) and combined control variables (*F=*2.69, *p=*0.117). All data: mean±SEM. \**p<*0.05; \*\**p<*0.01; \*\*\**p<*0.001.

**Extended Data Figure 5:**
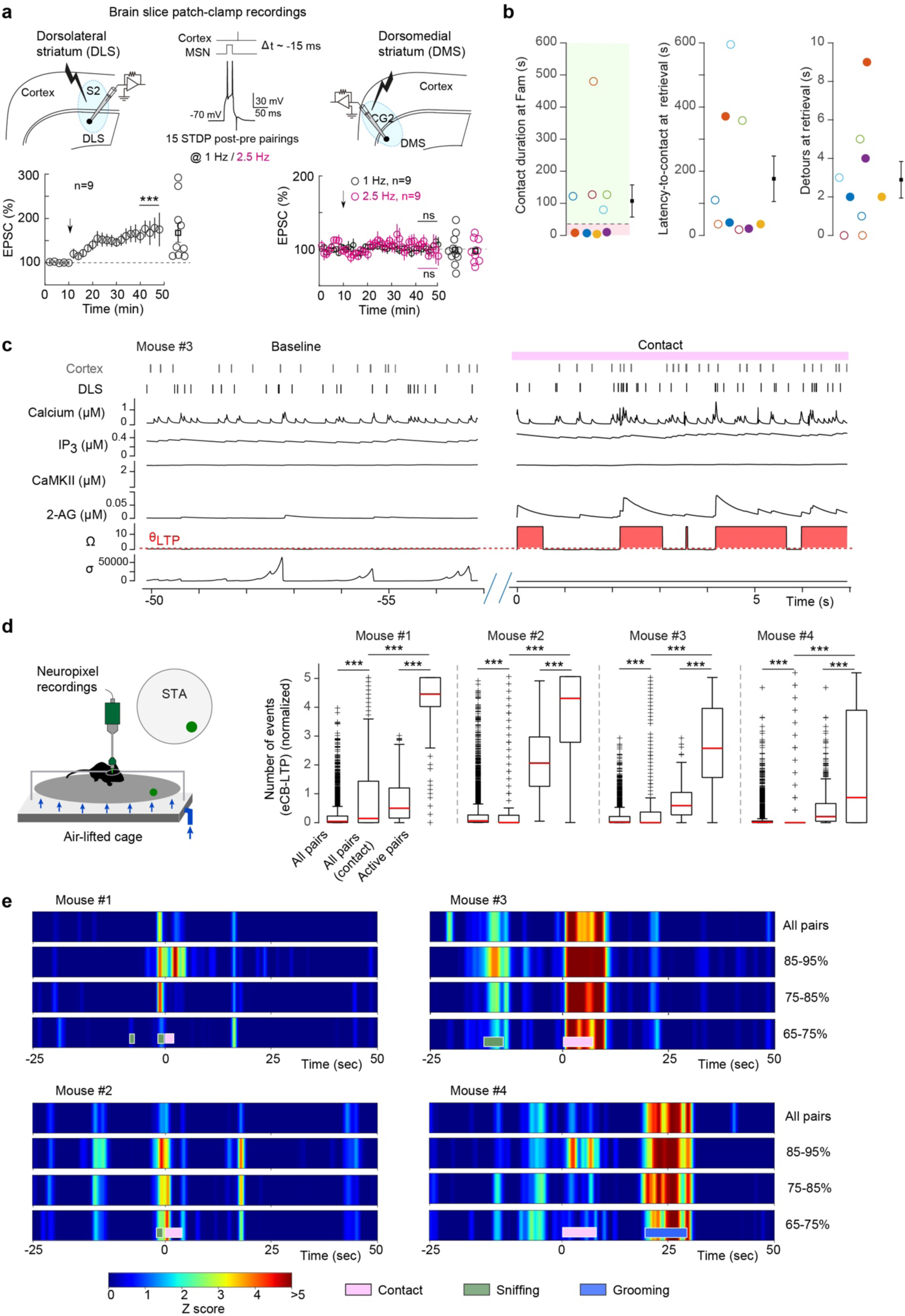
eCB-LTP expression in DLS and DMS, and eCB-LTP plasticitymeter. **a**, eCB-LTP is expressed in DLS but not in DMS. Top: Experimental set-up, with horizontal slices and parasagittal (with 30° angle) for DLS and DMS recordings, respectively. Bottom: Average time-courses of STDP. The arrow indicates the STDP pairing protocol. Scatter plots represent the percentage of change in EPSC amplitude 30-40 minutes after STDP pairings (one-sample t-test). Significant LTP was observed in DLS (*t=*3.1270, *p=*0.012, n=10 SPNs) but no significant plasticity was detected in DMS using pairings frequency at 1 or 2.5 Hz (*t=*-0.1397, *p=*0.892 at 1 Hz, n=9 SPNs; *t=*-0.408, *p=*0.694 at 2.5 Hz, n=9 SPNs). **b**, Contact duration at familiarization (<20 seconds: pink and >20 in green), latency-to-contact and detours at retrieval for mice in the MHC (n=9); filled circles: mice with contact duration <20 seconds (n=4 mice). **c**, Detailed evolution of calcium, IP_3_, active CamKII or 2-AG levels, and two variables associated to endocannabinoid-dependent plasticity, Ω and σ, for an illustrative spike train of a randomly-chosen active pair (mouse #3). eCB-LTP events arise (*red*) when Ω overcomes a threshold θ_LTP_ and σ is low (*Methods*), a situation that is more frequent during contact (*right*) rather than during baseline (*left*). **d**, Rate of eCB-LTP plasticitymeter events (events/second). The number of eCB-LTP events were preliminarily normalized for each cortico-striatal pairs by the sum of eCB-LTP events occurring during the baseline period. For each mouse (n=4), the average plasticimeter is very low when one considers all the possible cortico-striatal pairs over the whole experiment “all pairs” or during contact “all pairs (contacts)” (on average, 4853 pairs per experiment, ranging from 2646 to 8517 pairs). The average plasticitymeter increases when selecting the active pairs (*Methods*), especially during contact “active pairs (contact)”. Paired or independent two-sample t-tests, all showing *p*<0.001 between conditions for each mouse. **e**, Plasticitymeter z-scores across all pairs (top row) or pairs showing elevated correlated activity during contact show that the significant increase in eCB-LTP induction events during contact is robust to the choice of cortico-striatal pairs. The subset of pairs are selected based on the percentage of increase in correlated activity during contact relative to baseline, similarly to the *active pairs* (corresponding to the 95%-100% range, see main Fig. 3**,e** **c**),. Z-score maps are aligned to the start of contact. All data: mean±SEM. \**p<*0.05; \*\**p<*0.01; \*\*\**p<*0.001.

**Extended Data Figure 6:**
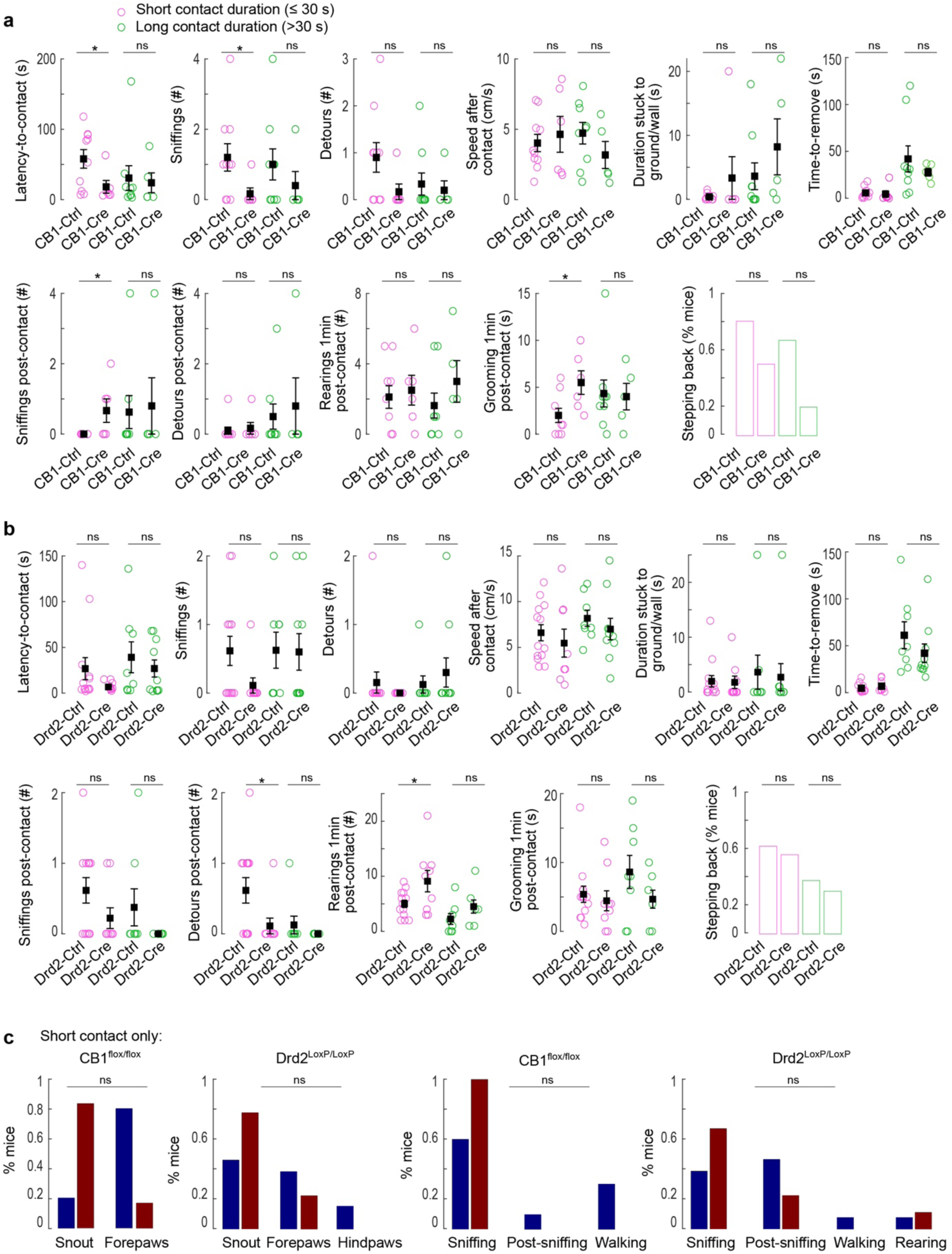
Behavioral characterization at familizarization of eCB-LTP conditional knock-out mice for presynaptic CB1R or D2R. **a**,**b**, Comparison of behavioral parameters at familiarization for CB1R^flox/flox^ mice (**a**) or *Drd2^LoxP/LoxP^* (**b**), clustered according to their contact duration. Statistical differences were observed between Cre and Ctrl mice with short contact duration with the sticky tape for a subset of parameters (two-sample independent t-tests). In CB1R^flox/flox^ mice: latency-to-contact (*p=*0.027); number of sniffings (*p=*0.049); number of sniffings post-contact (*p=*0.019); time spent grooming post-contact (*p=*0.034). In Drd2^LoxP/LoxP^ mice: number of detours post-contact (*p=*0.046), number of rearings post-contact (*p=*0.033). **c**, Comparison of the body part in contact with the sticky tape (left) or the activity immediately preceding contact (right) for mice having short contact duration. No statistical difference (log-linear regression model) was detected. All data: mean±SEM. \**p<*0.05; \*\**p<*0.01; \*\*\**p<*0.001.

**Extended Data Figure 7:**
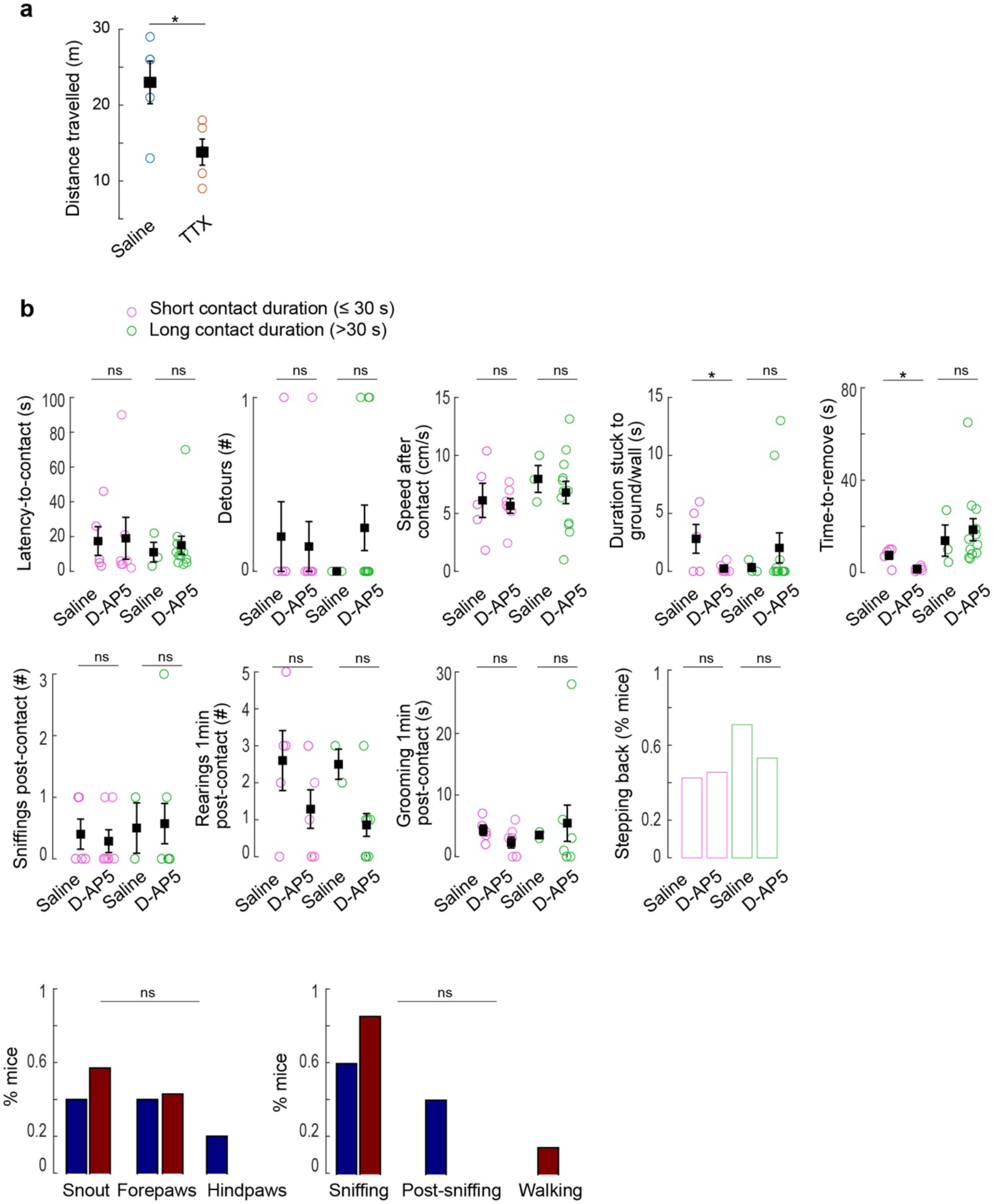
Control for D-AP5 absence of impact on retrieval performance. **a**, The distance travelled by mice injected with TTX (1 µM, 1-2 µL) was significantly lower than for mice injected with saline (2 µL) 45 min before open field test. Two-sample independent t-test: *t=*2.79; *p=*0.023. **b**, Comparison of the behavioral parameters at familiarization between mice injected with saline or D-AP5, clustered according to their contact duration. Significant differences were found between short contact duration for the following parameters (two-sample independent t-tests): duration of the mouse and tape stuck to ground/wall (*t=*2.45, *p=*0.034), time-to-remove (*t=*4.073, *p=*0.0022). All data: mean±SEM. \**p<*0.05; \*\**p<*0.01; \*\*\**p<*0.001.

**Extended Data Figure 8:**
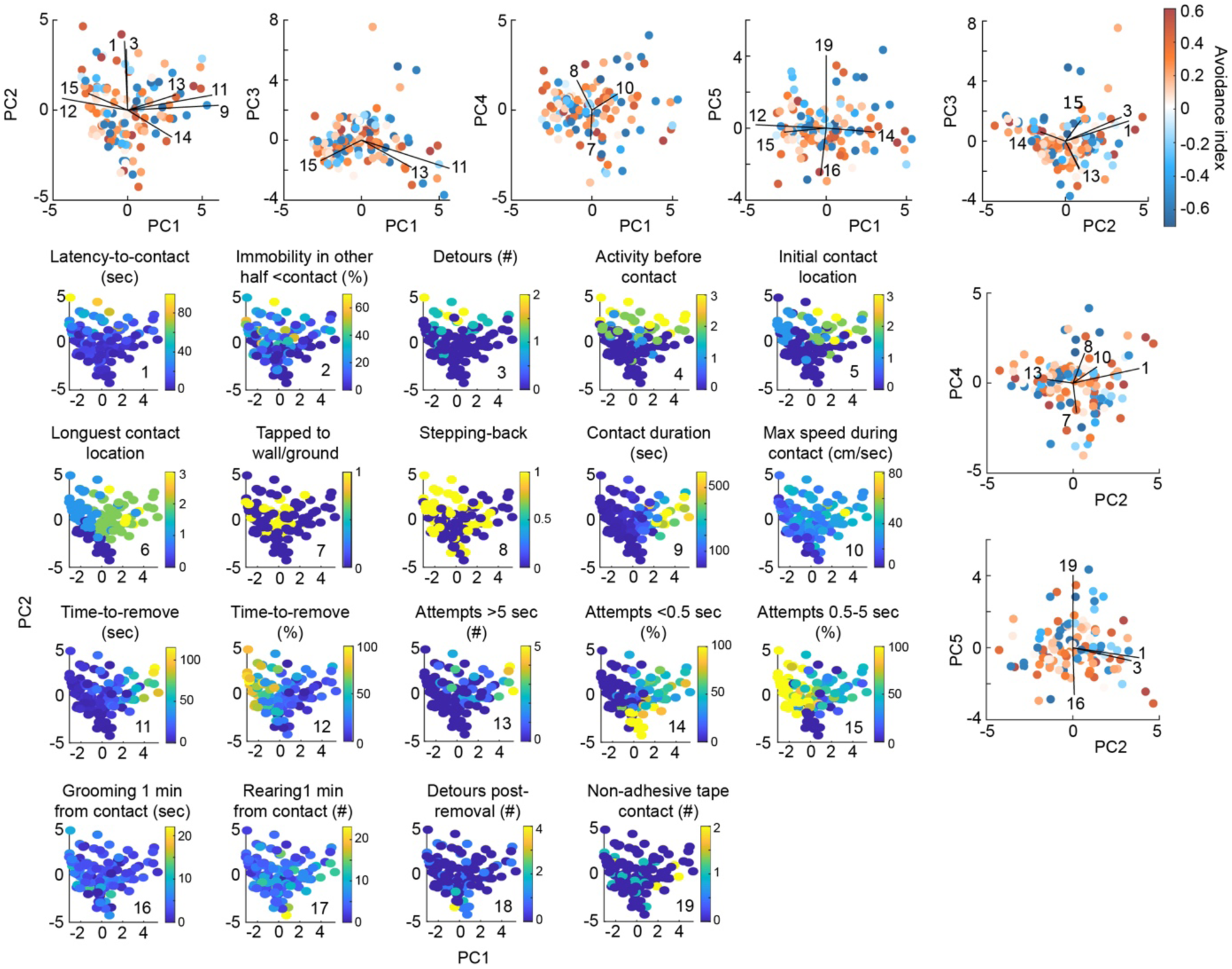
Impact of sticking features at familiarization on retrieval performance (related to the Supplementary Information) Binary comparison of mice according to avoidance index or number of detours estimated at retrieval for all mice (n=118, **a**) or in the subset of mice with contact duration ≤30 seconds (n=79 mice, **b**). Time spent grooming 1 min from contact (p<0.001) or 1 min post-removal for short contact (p<0.001) were significantly different (independent t-tests). No stepping back at contact and low surprise levels were associated to low numbers of detours (Chi-square tests: χ2=4.68, p=0.030 and χ2=10.8, p=0.013, respectively).

**Extended Data Figure 9:**
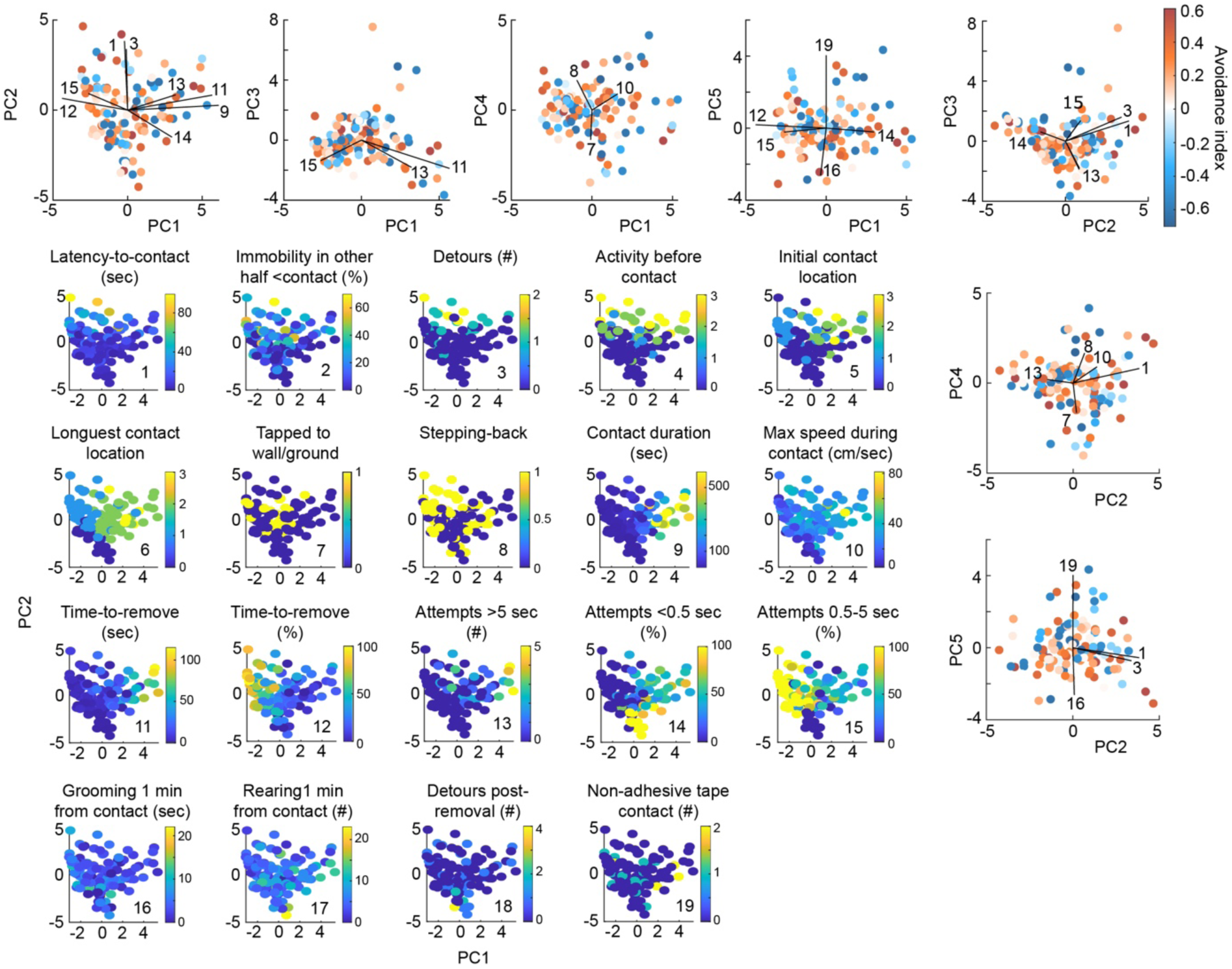
Multidimensional analysis of familiarization and impact on retrieval (related to the Supplementary Information) Projections of the 19 parameters considered in familiarization along the first two principal components (parula colormap). Projections of the avoidance index (blue to red colormap) along with the contributions of key parameters represented by vectors on the first five principal components.

**Extended Data Figure 10:**
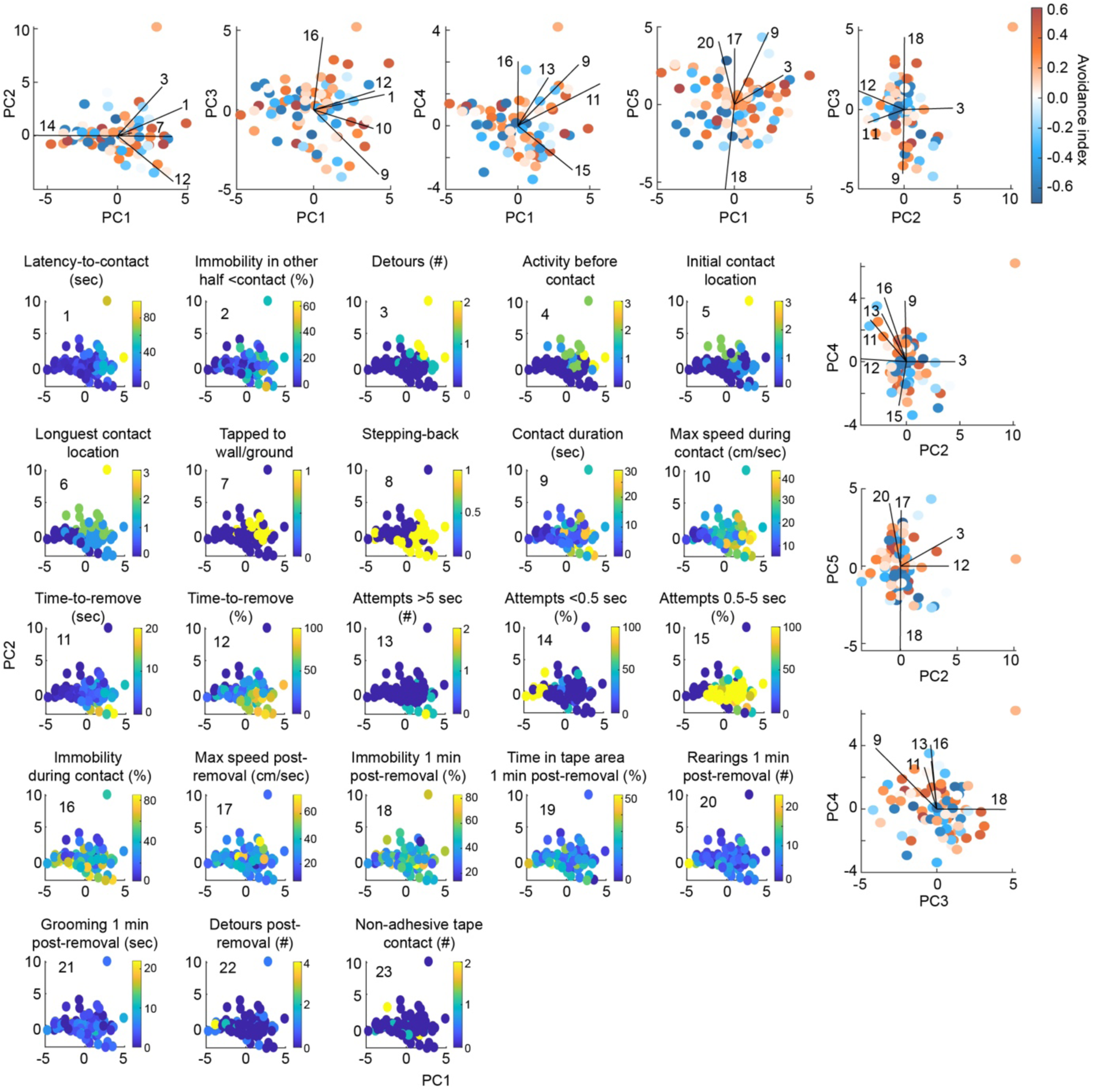
Multidimensional analysis of familiarization and impact on retrieval for mice exhibiting short contact duration with the sticky tape (related to the Supplementary Information) Projections of the 23 parameters considered in familiarization along the first two principal components (parula colormap) on mice exhibiting a short contact duration (≤30 seconds) with the sticky tape. Projections of the avoidance index (blue to red colormap) along with the contributions of key parameters represented by vectors on the first five principal components.

## Supplementary information

### Supplementary Methods

#### Object recognition memory: novel object recognition (NOR) task

The NOR task with four objects (Lego blocks) was run in a square Plexiglas arena (40x40cm) and consisted of three consecutive sessions at 24 hours intervals: (1) 10 min habituation to the empty arena, (2) 3 min familiarization to objects, and (3) 5 min retrieval session in which a novel object replaced one of the four objects. The object discrimination ratio was calculated as the ratio of the time spent exploring the novel object at retrieval over either (a) the average time spent on the four objects at familiarization or (b) the average time spent on the three familiar objects at retrieval.

### Procedural learning: accelerating rotarod

Mice were trained on the accelerating rotarod (Panlab, Barcelona, Spain) in 5 consecutive daily sessions of 5 trials (acceleration from 4 to 40rpm during 300 seconds). Between each trial, mice were placed in their homecage to rest for 15 minutes. Performance was assessed by subtracting the average latency of the last two trials of Day 1 (early state) or of Day 5 (late stage) to the average latency of the first two trials of Day 1.

### Reward-guided learning: cross-maze task

Mice access *ad libitum* to water containing 4% citric acid over one week. The north arm was used as starting point during the habituation/training period while entry to the south arm was blocked. During a 20 minutes habituation, the preferred west or east arm was determined by counting the number of visits. Then, the mouse was left another 20 minutes in the cross-maze with a cup of water at the extremity of the non-preferred arm (the baited arm). The training started on the next day and consisted of three consecutive daily sessions of 10 trials each, and the proportion of visits to the baited arm was quantified. The reward consisted of a drop of water (20μL). On the last day, a single probe test, in which the entry point of the mouse into the maze was shifted to the south arm (north arm was blocked) allowed to determine the response or place strategy of the mouse.

## Supplementary note

### Evaluation of the impact of sticking features at familiarization on retrieval performance

We aimed to detect if the behavior of the mouse during the first encounter with the sticky tape, at familiarization, could affect its behavior during retrieval. To do so, we first extracted parameters describing the sticking modalities and behaviors at familiarization, before, during and after contact with the sticky tape. We then performed multifactorial analyses and found that one-shot learning of STA could emerge regardless of the marked heterogeneity of behaviors recorded at familiarization.

We investigated a set of behavioral parameters larger than those briefly described in *Methods*. These are:

#### Period during the contact with the sticky tape

- the immediate post-sticking activity: stepping back {0: no stepping back, 1: stepping back}, average speed in the 5 seconds after contact (cm/s) and surprise level (scored from 0 to 3, based on mouse reaction, including stepping back, level of interruption in activity, unusual movements).
- latency before first active removal movement (seconds)
- number and percentage of epochs of voluntary movements aiming at tape removal; we classified them as short (<0.5 second), medium (0.5-1 second), long (1-5 seconds), or very long (>5 seconds) active attempts.
- time-to-remove (%): time-to-remove/contact duration
- taped to wall or ground: during contact, the tape could stick both to the mouse and to the ground or walls of the arena {0: <1 seconds, 1: ≥1 seconds}. The total duration spent taped to wall or ground was also considered.
- distance travelled (m)
- time spent immobile (%)
- average and maximal instantaneous speed (cm/s)
- rearings (#)
- time spent grooming (seconds)
- rearings (#) and time spent grooming (seconds) in the first minute following sticking, since 6% of mice did not successfully remove the tape.

#### Period after removal of the sticky-tape

- maximal speed in the ten seconds following sticky tape removal (cm/s)
- number of sniffings and detours (#)
- number of passages on the tape in the case when this one becomes stuck to the ground as a result of removal.

During the first minute following sticky tape removal (a preliminary comparison was made considering the first three minutes), we quantified:

- time spent immobile (%)
- time spent near the walls (%)
- time spent in the disk centered on the last position of the sticky tape and radius 8 cm (%)
- rearings (#)
- time spent grooming (seconds)

We first considered possible correlations between individual parameters evaluated at familiarization and the latency-to-contact, number of detours and avoidance index at retrieval (**Extended Data Figure 8**). We also performed independent t-tests using for binary classification, a avoidance index ≥0 or number of detours ≥2. We considered all mice or only those with short contact duration (≤30 seconds). We found that the time spent grooming (either 1 min from contact onset or 1 min post-removal) was significantly elevated at familiarization in mice that would subsequently show a clear avoidance response. In the short contact duration cases, we also noted that the percentage of mice that did not step back immediately after sticking to the tape was elevated in mice that showed less than two detours at retrieval (64 %, n=31 *vs*. 39%, n=48, Chi-squared test: χ^2^=4.68, *p*=0.030). In the same vein, lower surprise levels were found on average in mice showing less than two detours at retrieval (Chi-squared test: χ^2^=10.8, *p*=0.013).

To capture possible dependences between parameters and extract a contact configuration that would favor one-shot learning, we performed a multi-dimensional analysis (**Extended Data Figures 9 and 10**). We considered 19 parameters estimated at familiarization and valid for all mice: {1. latency-to-contact (s), 2. time immobile in other half before contact (%), 3. detours (#), 4. activity pre-contact, 5. first contact location (body part), 6. longest contact location, 7. tapped to the wall/ground (0/1), 8. stepping-back (0/1), 9. contact duration (seconds), 10. maximal speed during contact (cm/s), 11. time-to-remove (seconds), 12. time-to-remove (%), 13. active attempts >5 seconds to remove the tape (#), 14. attempts < 0.5 second (%), 15. attempts [0.5-5 seconds] (%), 16. grooming 1min after contact onset (seconds), 17. rearings 1 minute after contact onset (#), 18. detours post-removal (#), 19. non-sticky tape contacts post-removal (#)}. We then performed a factor analysis on mixed data (FAMD), using R-packages (*FactoMineR and factoextra*). Projections on the first two FAMD principal components of each individual parameter (**Extended Data Figure 9)**. We did not identify any clear hotspot from the mapping of the avoidance index onto the first FAMD principal components. A linear regression analysis was performed to extract dimensions that could predict the avoidance index at retrieval. When considering all mice on the first five dimensions (which explained 50.3% of the variance), it yielded a poor fit (r=0.252). A significant coefficient was found for the fifth principal component (*t*=-2.262, *p*=0.026). Grooming time 1 min after contact onset (contribution =15.39) and number of passages on the non-sticky tape (contribution =23.70) contributed the most to the fifth principal component.

We performed the same multifactorial analysis specifically on mice with short contact duration (≤30 seconds) (**Extended Data Figure 10**). We used 23 parameters (17/23 similar to the previous analysis): {1. latency-to-contact (seconds), 2. time immobile in the other half before contact (%), 3. detours (#), 4. activity pre-contact, 5. first contact location (body part), 6. longest contact location, 7. tapped to the wall/ground (0/1), 8. stepping-back (0/1), 9. contact duration (s), 10. maximal speed during contact (cm/s), 11. time-to-remove (seconds), 12. time-to-remove (%), 13. active attempts >5 seconds to remove the tape (#), 14. attempts <0.5 seconds (%), 15. attempts [0.5-5 seconds] (%), 16. time spent immobile during contact (%), 17. maximal speed in the first 10 seconds post-removal (cm/s), 18. time spent immobile 1 min post-removal (%), 19. time spent in scotch zone 1 min post-removal (%), 20. rearings 1 min post-removal (#), 21. grooming 1 min post-removal (seconds), 22. detours post-removal (#), 23. non-sticky tape contact post-removal (#)}. The linear regression performed on the first five dimensions (explaining 50.2% of the variance) poorly predicted the avoidance index at retrieval (r=0.266), with no significant regression coefficients.

We tested non-linear classification algorithms either directly on 23 parameters of familiarization or on the first five FAMD components. The classification boundary was set according to the avoider rate boundary (avoidance index >0). k-nearest neighbors (k-NN, with k=5) and decision trees were trained on 75% of the dataset (n=79 mice) and cross-validated (reaching 67-85% accuracy for k-NN and 90-95% for decision trees). Yet, classifiers performed at chance level on the test data set. We also tested other classification or regression indices (number of detours, latency-to-contact), yielding poorer results on the cross-validation dataset, and remaining at chance level on the test data set.

To conclude, this extensive analysis of mouse behavior at familiarization did not reveal the existence of key variables predicting subsequent learning and avoidance behavior on retrieval.

